# *Chlamydia*-driven ISG15 expression dampens the immune response of epithelial cells independently of ISGylation

**DOI:** 10.1101/2024.05.27.596023

**Authors:** Yongzheng Wu, Chang Liu, Chongfa Tang, Béatrice Niragire, Yaël Levy-Zauberman, Cindy Adapen, Thomas Vernay, Juliette Hugueny, Véronique Baud, Agathe Subtil

## Abstract

Excessive inflammation upon *C. trachomatis* infection can cause severe damages in the female genital tract. This obligate intracellular bacterium develops mainly in epithelial cells, whose innate response contributes to the overall inflammatory response to infection. The ubiquitin-like protein interferon-stimulated gene 15 (ISG15) stimulates interferon γ (IFNγ) production and is required for bacterial clearance in several infectious contexts. Here, we describe and investigate the consequences of the increase in ISG15 expression by epithelial cells infected with *C. trachomatis*. Infection of HeLa cells and primary ecto-cervical epithelial cells resulted in a transcriptional up-regulation of *ISG15* expression. This did not involve the canonical IFN-I signaling pathway and depended instead on the activation of the STING/TBK1/IRF3 pathway. Absence or reduction of ISG15 synthesis led to increased production of several cytokines and chemokines including interleukin (IL) 6 and IL8, implicating that ISG15 normally dampens the immune response induced by *C. trachomatis* infection in epithelial cells. ISG15 exerted its control from an intracellular location, but without involving ISGylation. Finally, higher levels of inflammation and delayed bacterial clearance were observed in the genital tracts of ISG15-KO mice infected by *C. trachomatis* compared to wild type animals, however IFNγ production was unchanged. Altogether, our data show that ISG15 expression acts as a brake on the immune response to *C. trachomatis* infection in epithelial cells and limits bacterial burden and inflammation in mice.

## Introduction

Mammalian cells possess various sensors to detect invading microorganisms and activate innate defense mechanisms, especially the secretion of inflammatory cytokines and chemokines, aimed at the elimination of the intruders. Interferon stimulated gene 15 (ISG15) is a protein of 15 kDa with two ubiquitin-like domains. Similar to ubiquitin, it can be conjugated to a lysine residue on a target protein through a series of enzymatic reactions involving E1/E2/E3 ligases. ISG15 conjugation (ISGylation) is rendered possible by the LRLRGG motif exposed after the removal of eight amino acids at the C-terminus of the protein (1). ISG15 also displays biological activity as a free molecule, from two locations: it can be secreted and exert its activity through binding to a specific membrane receptor or it can act as a free cytoplasmic protein, by unknown mechanism(s) (2–4). ISG15 and the members of the enzymatic cascade that lead to ISGylation are strongly induced by type I interferon (IFN-I) and ISG15 was initially studied for its anti-viral property in mice (5). However, against expectations, inherited ISG15-deficient patients do not show increased susceptibility to viral infection. In contrast, weakly virulent mycobacteria such as BCG, which only triggers a mild inflammatory response in normal individuals, induces a life-threatening infection in patients lacking ISG15 (6). It was shown that lack of ISG15-dependent induction of IFNγ was responsible for the severe consequences of ISG15 deficiency in humans (6). More recently, it was reported that ISG15 expression was induced independently of IFN-I upon infection of epithelial cells by a gram-positive bacterium, that developed in the cytoplasm, *Listeria monocytogenes* (7). Like in the case of *Mycobacterium*, ISG15 expression restricted *Listeria* development, supporting the idea that this ubiquitin-like molecule was exerting anti-bacterial activity(ies).

*Chlamydia trachomatis* is an interesting infectious bacterium to explore further the role of ISG15 in the innate host response to bacterial infection, since it is a gram-negative bacterium, that develops in a different cell type than *Mycobacterium*, and in a vacuole, unlike *Listeria*. *C. trachomatis* is the leading cause of bacterial sexually transmitted infections (STIs) in the world and of preventable blindness of bacterial origin, and constitute one major problem for public health worldwide (8). This obligate intracellular bacterium grows exclusively inside a vacuolar compartment, the inclusion, within epithelial cells of genital or conjunctival mucosae. Genital *C. trachomatis* infections are very common, with a worldwide estimate of more than 100 million new cases every year (9). Prevalence is difficult to record because most cases are asymptomatic and go undetected. Chronic and/or repeated infection can eventually cause severe and irreversible consequences, particularly in women, such as pelvic inflammatory disease and scar formation in the fallopian tubes leading to tubal infertility. While it appears that the tissue damages are due to inadequate levels of immune response of the host, the conditions that favor these deleterious consequences are still poorly understood.

The importance of innate immune response against *Chlamydia* infection has been highlighted by the finding that innate host defense is sufficient for eradicating *C. trachomatis* from the FGT of mice(10). As other microorganisms, *C. trachomatis* have acquired several mechanisms to subvert host defense (11). For instance, the bacteria express a protease CPAF (*Chlamydia* protease-like activity factor) that is capable of blocking the secretion of neutrophil extracellular traps by neutrophils (12), of degrading host antimicrobial peptides (13), and, possibly indirectly, of inhibiting NF-κB p65 translocation to the nucleus (14). The bacteria also secrete a deubiquitinase that inhibits ubiquitination primed by IFNγ in human epithelial cells, thereby evading subsequent elimination by the host (15). TepP, an effector of *C. trachomatis* secreted at early stage of infection, can also manipulate the host signals associated with innate immunity of host cells (16, 17).

Low- and high-risk human papilloma virus (HPV) infection elicits differential ISG15 induction in the cervical mucosae of infected human patients, indicating that ISG15 may contribute to the innate response to infection in the FGT (18). Furthermore, given the key role of IFNγ in resolving *C. trachomatis* infection (19–21) and the fact that ISG15 is described as a potent IFNγ-inducing cytokine playing an essential role in antimycobacterial immunity (6), we anticipated that ISG15 might be an important player for efficient clearing of *C. trachomatis* infection. Indeed, we observed a strong induction of ISG15 expression by epithelial cells upon infection by *C. trachomatis*. We uncovered the signaling cascade that led to an increase in *ISG15* transcription and studied its outcome on infection.

## Methods

### Cells

The human cervical epithelial HeLa cell line was from ATCC (ATCC^®^CCL-2™). The cells were grown and propagated in Gibco^®^ Dulbecco’s Modified Eagle Medium (DMEM) with GlutaMAX (Thermo Fisher Scientific) and 10% of fetal calf serum (FCS) at 37 °C with 5% of CO_2_. Primary ecto-cervical epithelial cells were isolated as described previously(22) and cultured in Gibco® keratinocyte serum-free medium (K-SFM) supplemented with recombinant human EGF and bovine pituitary extract (Thermo Fisher Scientific, #17005075). The cells were propagated in 75cm^2^ flasks (Falcon) and seeded in culture dish (0.1×10^6^ cell/well for 24-well plates or 0.2×10^6^ cell/well for 12-well plates) 24 h before infection or treatment.

ISG15-KO HeLa cells were obtained following a standard procedure(23). Briefly, 0.5 μg of pSpCas9(BB)-2A-Puro (PX459) V2.0 plasmid (Addgene #62988) containing guide RNA (gRNA) duplex sequence of human ISG15 (F: 5’-ggcgtgcacgccgatcttct; R: 5’-agaagatcggcgtgcacgcc) inserted with BbsI enzyme, was transfected into HeLa cells (0.15 × 10^6^ cells/well, 24-well plate) seeded overnight prior to transfection using 1 μl of jetPRIM® (Polyplus). Four hours later, medium was replaced by fresh medium containing puromycin (2 μg/ml) and cells were incubated for 24h. The surviving cells were then detached, counted, and serially diluted to collect individual clones in 96-well plates. Absence of ISG15 expression was verified on individual clones by western blot.

### Bacteria

*C. trachomatis* serovar LGV-L2 (434/Bu) and LGV-L2^IncD^GFP bacteria expressing constitutively GFP(24) were purified on a density gradient as previously described(25). In brief, 80% confluent HeLa cells were infected with LGV-L2 (MOI=1) in the presence of cycloheximide (1 μg/ml). Forty-eight hours later, the cells were homogenized mechanically with sterile glass beads followed by sonication to release bacteria. Infectious bacteria were purified by density gradient centrifugation using Gastrografin^®^ (Bayer, Germany), resuspended in sucrose-phosphate-glutamic acid (SPG) buffer (10 mM sodium phosphate [8 mM Na_2_HPO_4_-2 mM NaH_2_PO_4_], 220 mM sucrose, 0.50 mM l-glutamic acid), aliquoted and stocked at -80°C.

### siRNA treatment

siRNA smartpools were purchased from Dharmacon (Lafayette, USA) except for siRNAs for ISG15 which were individually purchased from Eurogentec (Seraing, Belgium). One and half microliter siRNA (10 μM) was mixed with 0.5 μl Lipofectamine iMax (Invitrogen) in 50 μl of Opti-MEM™ media (Gibco®) for 10 min at room temperature according to the manufacture recommendation. The mixture was added into a culture dish before adding cells (0.1×10^6^ cells/well for 24-well plate) suspended in 0.45 ml DMEM/FCS medium (final concentration of siRNA was 30 nM), mixed and incubated for 48 h before treatment or infection. The siISG15 oligos were mixed (10 μM each) #1 5′-uccuggugaggaauaacaa dTdT-3′; #2 5′-gcaccguguucaugaaucu dTdT-3′)(26).

### Cell infection

Cells, seeded the previous day in culture dish (0.1×10^6^ cell/well for 24-well plate or 0.2×10^6^ cell/well for 12-well plate), were rinsed twice using pre-warmed media before adding LGV-L2 in the culture media. Twenty-four to 42 h later, the culture medium was collected, spined for 15 min at 1,500 ×*g* to remove cell debris, frozen at -20 °C and used for later cytokine assay by ELISA. The cells were lysed either in urea buffer (8 M urea, 30 mM Tris, 150 mM NaCl, 1% v/v sodium dodecyl-sulfate pH 8.0) for protein extraction or in RLT buffer (Qiagen) for RNA extraction. In certain experiment, the cells were pre-treated with pharmacological inhibitors for 30 min or transfected with siRNA for 48 h before bacterial infection. Wortmannin was from Selleckchem (#S2758).

### Progeny assay

Cells (0.1×10^6^ cells/well) plated the day before in 24-well plates were infected with LGV-L2^IncD^GFP bacteria expressing constitutively GFP (24) at a MOI = 0.15. Forty-eight hpi, cells were detached, lysed using glass beads and the supernatant was used to infect HeLa cells plated the day before (0.1×10^6^ cells/well in a 24-well plate), in serial dilutions. Twenty-four hours later, cells with an infection lower than 30 % (checked by microscopy) were detached using 0.5 mM EDTA in PBS and fixed in 4% (w/v) paraformaldehyde (PFA) 4% (w/v) sucrose in PBS, before analysis by CytoFLEX flow cytometer (Beckman Coulter). The data were analyzed using FlowJo (version 10.0.7) to determine the bacterial titer.

### Bacterial adhesion and entry studies

*C. trachomatis* binding to or entry into epithelial cells was performed as described previously (24). For binding experiments, pre-chilled cells (30 min at 4 °C) grown on coverslips were incubated with LGV-L2^IncD^GFP (MOI = 30) for 4 h at 4 °C (Bacteria were gently sonicated before infection to disrupt bacterial aggregates). Cells were washed 3 times with chilled DMEM medium, the cells were fixed with 4% (w/v) paraformaldehyde (PFA) and 4% (w/v) sucrose for 20 min at room temperature permeabilized by 0.05% saponin/PBS/0.1% BSA solution. DNA was stained by Hoechst (0.5 μg/ml) before manually counting GFP-bacteria per cell in several fields taken with a Deltavision™ microscope (GE, UK). For bacterial entry study, pre-chilled cells seeded on coverslip were incubated with LGV-L2^IncD^GFP (MOI = 10) for 45 min at 4°C. The cells were then washed with pre-warmed medium for 3 times and incubated at 37 °C for the indicated time before fixation as above. Extracellular bacteria were stained with a mouse anti-MOMP-LPS antibody (Argene #11–114) followed with Cy5-conjugated anti-mouse secondary antibody (Amersham Biosciences). The bacterial entry was presented as ratio between intracellular bacteria (GFP only dots) and total bacteria (all GFP positive dots including GFP/Cy5 dots).

### Immunofluorescence

Cells (0.1×10^6^ cells/well) seeded on coverslips in a 24-well culture dish were infected with LGV-L2 strain with MOI = 1 for 40 h or treated with IFNα (Biogen Idec, USA) for 24 h. The cells were then fixed in PFA as described above. After fixation, the cells were washed in PBS and incubated with 50 mM NH_4_Cl to quench the residual PFA, followed with 10 min in 0.05% (w/v) saponin, 0.1% (w/v) BSA for cell permeabilization. Intracellular bacteria and ISG15 were stained with a mouse anti-MOMP-LPS (Argene # 11-114) antibody and a rabbit anti-human ISG15 antibody (a kind gift from E. C. Borden, Cleveland Clinic, Cleveland, OH, USA) followed with Alexa488-conjugated anti-mouse and Cy5-conjugated anti-rabbit secondary antibody. The dilutions were made in PBS containing 1 % of BSA and 0.05 % saponin. DNA was stained using 0.5 µg/ml of Hoechst 33342 (Thermo Fisher Scientific) added in the secondary antibody solution. Coverslips were mounted on slides in a Mowiol solution. Images were acquired on an Axio observer Z1 microscope equipped with an ApoTome module (Zeiss, Germany) and a 63× Apochromat lens. Images were taken with an ORCA-flash4.0 LT camera (Hamamatsu, Japan) using the software Zen. The images of ISG15-KO cells complemented with *ISG15* gene using sleeping-beauty system were directly observed under an Axio Observer microscope equipped with a filter to record GFP fluorescence (Zeiss, Germany). TNFα used in in Fig. S5 was from R&D system (#210-TA-005/CF).

### Complementation of ISG15-KO cells by constitutively over-expressing ISG15

ISG15 sequence with full-length or depletion of 6 nucleotides encoding 2 glycine residues at the C-terminus of the protein was cloned into pSBbi-GP plasmid (addgene, #60511) by Gibson assembly and verified by sequencing. In this plasmid, the same constitutive promoter controls expression of GFP and ISG15. Plasmids were transfected into ISG15-KO cells using jetPRIME transfection reagent (Polyplus) and the culture medium was replaced 4 h later. Twenty-four hours post transfection, the transfected cells were selected by puromycin (1 μg/ml). The medium containing puromycin was replaced every 2 days to remove dead cells. Selection efficiency was evaluated under the microscope by monitoring GFP expression. Seven to 10 days post puromycin selection, almost all cells were GFP-positive and cell pools were frozen or kept in culture for the experiments.

### RT-PCR and quantitative PCR

Total RNAs were isolated from infected or treated cells using RNeasy Mini Kit (Qiagen) and reverse transcription (RT) was performed using the M-MLV Reverse Transcriptase (Promega). The genomic DNA of mouse tissue was extracted using DNeasy Blood & Tissue Kit (Qiagen). Briefly, the frozen upper FGT was homogenized in pre-chilled Lysing Matrix D 2 mL tube (MP Biomedicals) containing 400 μl PBS using Precellys 24 tissue homogenizer (Bertin, Germany). After pelleting the tissue debris (14000 ×*g*, 2 min), an aliquot of the supernatant (200 μl) was used to extract genomic DNA. The qPCR was conducted on the complementary DNA (cDNA) or genomic DNA with LightCycler 480 system (Roche) using SYBR Green Master I (Roche). For bacteria load detection, serial dilutions of known quantities of 16S DNA of *C. trachomatis* and mouse β-actin DNA were used as standardized templates for qPCR. Data were analyzed using the 2^-ΔΔCt^ method and results were presented as Log2 fold changes compared to uninfected control(27), or as ratio of *Chlamydia* 16S (pg) to host DNA (ng) for the bacterial load. The primers used for qPCR are described in Table S1.

### Western blot, dot blot, EMSA and ELISA assays

Equal volumes of cell lysates were subjected to SDS-PAGE, transferred to polyvinylidene difluoride (PVDF) membranes and immunoblotted with the proper primary antibodies diluted in 1× PBS containing 5% milk and 0.01% Tween-20. Primary antibodies used were rabbit anti-human ISG15, rabbit anti-heat shock protein 60 of *C. trachomatis* (in house preparation), rabbit anti-Akt (Cell Signaling, #4691) and anti-p-AKT antibodies (Cell Signaling, #4060), mouse anti-human β-actin (Sigma, #A5441). Immunoblots were analyzed using horseradish peroxidase secondary antibodies, and chemiluminescence was analyzed on an Amersham ImageQuant™ 800 imaging system (Cytiva, USA).

To assess ISG15 secretion by HeLa cells, one millimeter of cell-free culture medium collected from 12-well plate was loaded by several times to a PVDF membrane using the Convertible Filtration Manifold System (Gibco-BRL, # 11055) followed with immunobloting as described above. Recombinant human ISG15 (R&D systems, #UL-601-500) resuspended in culture medium served as a positive control in this experiment.

Electrophoretic mobility shift assay (EMSA) was performed using the HIV-LTR tandem κB oligonucleotide as a κB probe (28).

The human XL cytokine luminex® performance assay 44-plex fixed panel (R&D systems, #LKTM014) was used to measure cytokine secretion in the culture medium. The culture supernatant was spined and diluted 5 times in fresh medium, a 50 μl aliquot was used for ELISA assay following the manufacturer instructions. To determine cytokine/chemokine concentrations in the genital tracts of mice, the homogenate of mouse tissue was prepared as described above. Non-diluted supernatant of tissue homogenate (50 μl) was applied to Bio-plex pro mouse cytokine & chemokine assay kit (Bio-Rad, #M60-009RDPD).

### Single cell cytokine detection by flow cytometry

Cells (0.2×10^6^ cells/well) plated the day before in 12-well plates were infected with LGV-L2^IncD^GFP bacteria expressing constitutively GFP(24) at a MOI = 1. Twenty-four hpi, the cells were incubated with 5 μg/ml brefeldin A (Biolegend) for 6h to block cytokine secretion. Cells were then detached using 0.5 mM EDTA in PBS and fixed in 4% (w/v) paraformaldehyde (PFA) 4% (w/v) sucrose in PBS, before permeabilization with 0.3% (v/v) Triton X-100 in PBS for 10 min and block in 1% BSA in PBS for 1h. The cells were then incubated 0.1 % BSA in PBS with or without 1/40 dilution of PE/Cyanine7-conjugated anti-human IL6 (Biolegend, #501119) or 1/80 dilution of PE-conjugated anti-human IL8 (Biolegend, #511408) for 1h. After washes, the cells were resuspended in PBS and single cell fluorescence was measured on a CytoFLEX flow cytometer (Beckman Coulter). Cells not stained or stained with single color were used for fluorescence compensation. The data were analyzed using FlowJo (version 10.0.7) and the data from the IL6^+^/GFP^+^ or IL8^+^/GFP^+^ populations were exported with scale values for further analyzed by Prism9 (GraphPad).

### Mice and infection

Female c57BL/6J mice were purchased from Charles River Laboratories (France). ISG15-KO mice, also in the c57BL/6J background, were kindly provided by Dr Klaus-Peter Knobeloch (Leibniz-Institut für Molekulare Pharmakologie (FMP), Germany). The animals were 6-7 weeks old. All animals were given 2.5 mg of medroxyprogesterone (Depo-provera®, Pfizer) s.c. 7 days prior to infection to synchronize the menstrual cycle. The experiments were performed in accordance with the French national and European laws regarding the protection of animals used for experimental and other scientific purposes. The study was approved by the ethics committee of Institut Pasteur (approval N° 2014-0054). The animals were anesthetized by intraperitoneal injection of Ketamine/Xylazine suspended in PBS. *C. trachomatis* (10^6^ IFU/mouse) suspended in 5 μl of SPG was introduced directly to the uterine horn through non-invasive trans-cervical injection using NSET® device (ParaTechs Cat # 60010). This procedure was previously shown to induce chronic infection in the upper FGT of mice(29). The same volume of SPG was injected in the upper FGT of control animals. At the indicated times post infection, the animals were sacrificed by cervical dislocation. The upper genital tracts (uterine horn, fallopian tubes and ovaries) were excised and frozen immediately into liquid nitrogen. For morphological changes, 4 weeks post infection, the excised FGTs were rinsed in PBS and imaged using a Nikon camera (D700 with the Micro-Nikkor 105 mm objective). The hydrosalpinx score of the uterine horn was quantified as described (30).

### Statistical analysis

Data were analyzed using Prism9 (GraphPad). Non-paired t-tests, paired t-tests and one-way ANOVA with post-hoc Tukey tests were used for two group and multiple group comparisons, respectively. The correlation between cytokines and bacterial loads was determined with two-tailed Pearson test.

## Results

### Chlamydia infection enhances ISG15 expression in epithelial cells

ISG15 expression during *C. trachomatis* infection was first examined in HeLa cells, a cancer cell line derived from the cervical epithelium. Infection of HeLa cells with *C. trachomatis* serovar L2 (LGV-L2) led to an increase in ISG15 levels in a time- and dose-dependent manner (Fig. 1A-B), which correlated with an increase in *ISG15* transcripts 6 hours post infection (hpi) and onwards (Fig. 1C). The increase in ISG15 levels in infected cells was also detected by immunofluorescence (Fig. 1D). Comparison with IFNα and IFNβ, well-established inducers of ISG15 transcription, showed that, although robust, ISG15 levels upon infection were lower than upon exposure to 10 ng/ml IFN-I (Fig. 1E). An increase in *ISG15* transcription and higher ISG15 protein levels were also observed in primary cervical epithelial cells isolated from patient explants (Fig. 1F). We concluded from these observations that ISG15 synthesis by epithelial cell is increased upon infection by *C. trachomatis*.

**Fig 1.**
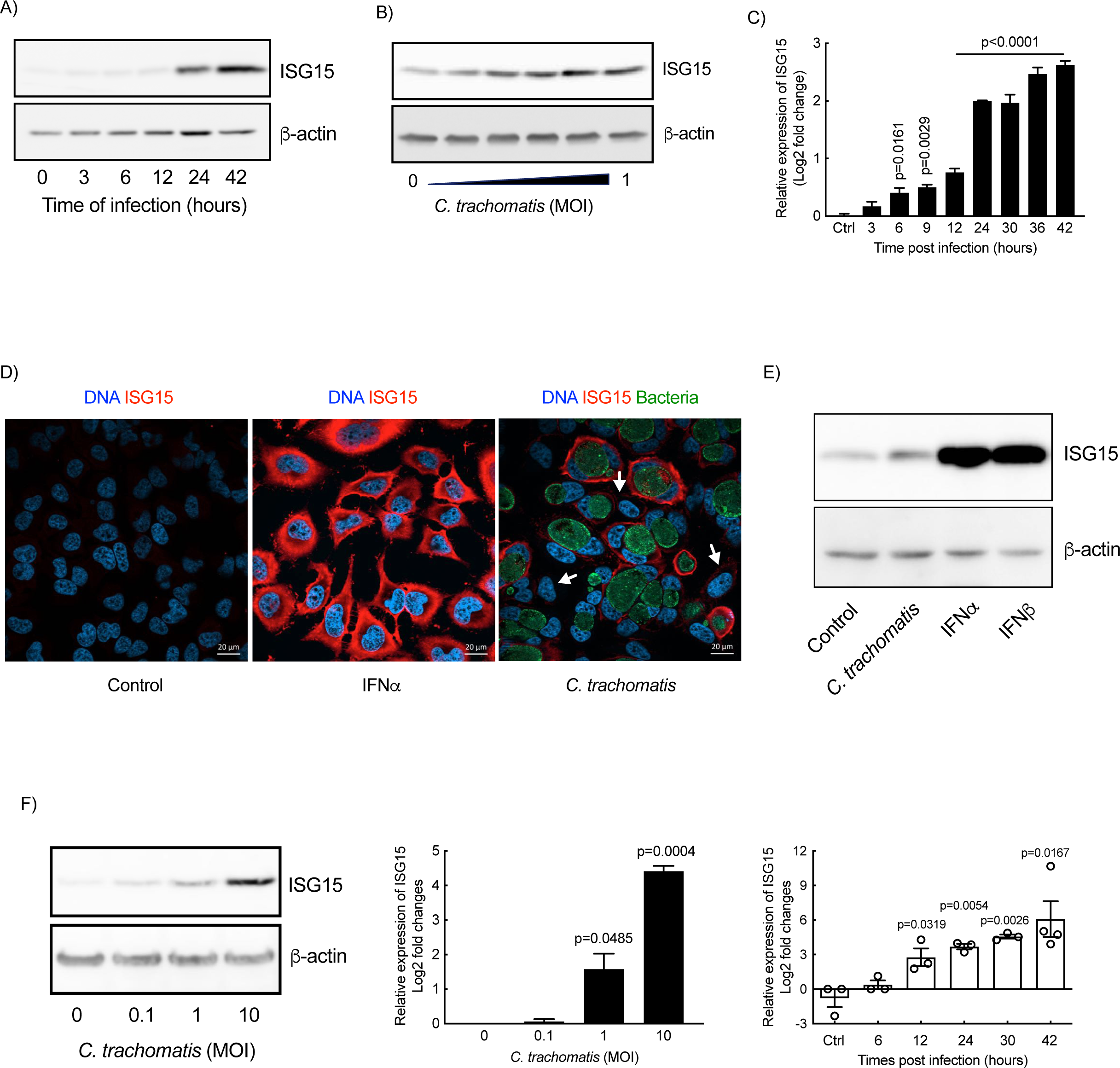
*C. trachomatis* infection induces ISG15 synthesis in epithelial cells. A-C) Time-course of infection of HeLa cells with *C. trachomatis* (A and C, MOI=1) or with increasing MOI (B, 42 hpi). After infection, ISG15 transcript (C) and protein (A-B) levels were examined by RT-qPCR and western blot, respectively. D) HeLa cells were incubated with IFNα (10 ng/ml) or *C. trachomatis* (MOI=1) for 30 h followed by immunostaining. E) HeLa cells were incubated with *C. trachomatis* (MOI=1) for 42 h or IFN-I (10 ng/ml) for 24 h before determining the expression of ISG15. F) Primary epithelial cells were incubated with *C. trachomatis* at the indicated MOI. ISG15 expression was examined 42 h post infection (left and middle panels) or at the indicated times post infection (right panel). The western blots and immunofluorescence data are representative of at least 2 independent experiments. All other data represent 3 independent experiments. One-way ANOVA test (time course) and unpaired t-test (dose course in primary cells) were performed, and the p-value of the comparison with uninfected control is shown.

### ISG15 dampens C. trachomatis-induced immune response

One of the documented consequences of *C. trachomatis* infection in epithelial cells is the production of the pro-inflammatory cytokines IL6 and IL8 (31, 32). ISG15 is a negative regulator of IFN-I immunity, preventing over-amplification of inflammation (4). The observation that ISG15 was strongly induced during *Chlamydia* infection prompted us to test whether it modulated *Chlamydia*-induced inflammation. To this end, we silenced *ISG15* expression in HeLa cells using small interfering RNA, which resulted in a strong reduction in ISG15 level, below detectable level (Fig. 2A, left panel). We observed an increase in the transcription levels of the two pro-inflammatory cytokine genes *IL6* and *IL8* upon silencing of *ISG15* (Fig. 2A, top panels). This was correlated with an increase in the amount of cytokine in the culture supernatant (Fig. 2A, bottom panels). In the rest of the manuscript, only transcriptional data are shown, as we consistently observed a good correlation between the two read-outs. We next used murine embryotic fibroblasts (MEFs) lacking ISG15 (7) to measure the innate response to infection. Consistent with what we had observed in human cells, mouse cells KO for *ISG15* displayed increased synthesis of IL6, and to a lesser extent of IL8 (KC), upon *Chlamydia* infection (Fig. 2B). Finally, we disrupted *ISG15* expression in HeLa cell using CRISPR/Cas9 and obtained four clones homozygotes for *ISG15*, with a disruption of the open reading frame resulting in the absence of ISG15 expression (Fig. S1A). In three of them we observed an increase in *IL6* and *IL8* transcription upon infection, compared to the parental wild-type cell line (Fig. S1B). This observation confirms that overall disruption of *ISG15* expression increases the expression of *IL6* and *IL8*, but that compensatory mechanisms may take place in some clones. Two clones C2 and C4 were selected for complementation experiments using the sleeping-beauty transposon system to achieve expression in the whole cell population (Fig. 2C, left panel; S1C)(33). Constitutive expression of ISG15 in C2 and C4 reduced basal and *Chlamydia*-induced transcription of *IL6*/*IL8* (Fig. 2C). Multiplex ELISA assay on culture supernatants confirmed infection-stimulated secretion of IL6 and IL8, as well as of several other cytokines and chemokines, was reduced in cells expressing ISG15 compared to ISG15-KO HeLa cells (Fig. S2). The ISG15-sensitive secretome included pro-inflammatory cytokines/chemokines, anti-inflammatory cytokines such as IL10 and IL13, and other actors of the immune response (*e.g*. TNFSF5, TNFSF10, FLT3L, granzyme B, etc). Altogether, these data show that ISG15 normally acts as a break on the host immune response to *C. trachomatis* infection.

**Fig 2.**
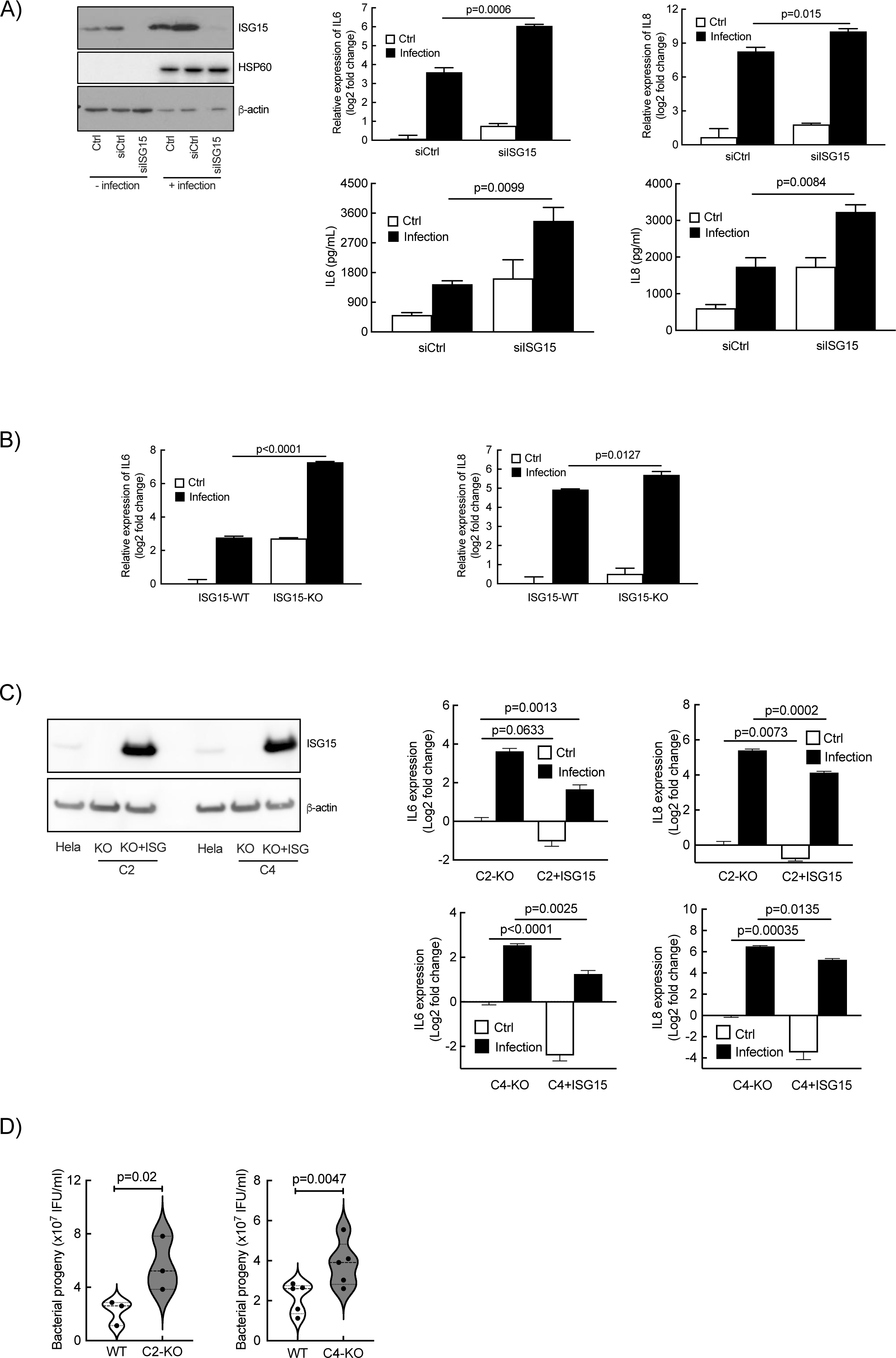
ISG15 exerts a negative control on *IL6* and *IL8* expression. A) Cells were transfected with siRNA targeting human *ISG15* or non-relevant oligonucleotides for 48 h prior to infection with *C. trachomatis* (MOI=1). Cells were collected 42 hpi. ISG15 levels were determined by western blot (left panel). HSP60, a bacterial protein, control for bacterial load, and actin serves as a loading control. The *IL6* and *IL8* transcript (top histograms) and protein (bottom histograms) levels were determined by RT-qPCR and ELISA, respectively. B) MEFs lacking ISG15 or not were infected with *C. trachomatis* (MOI=1) and the amount of *IL6* and mouse *IL8* (KC) transcripts were determined 42 hpi by RT-qPCR. C) *ISG15*-deficient clones (C2 and C4) were complemented with ISG15 (left panel) and infected with *C. trachomatis* (MOI=1). Transcripts levels for *IL6* and *IL8* were measured 42 hpi (middle and right panels). D) Quantification of infectious bacteria collected in the indicated cellular backgrounds 48 hpi. Each dot in panel D represents an individual experiment. The rest of the data are representative of at least three independent experiments. Unpaired t test was conducted in A and B, one-way ANOVA was performed in C, and paired t-test was used in D.

#### ISG15 expression restricts C. trachomatis development

We next investigated whether the absence of ISG15 expression affected *C. trachomatis* development. Wild-type and ISG15-KO HeLa cells were infected with *C. trachomatis* for 48 h (a duration corresponding on average to the completion of one developmental cycle) before measuring the progeny. We observed a moderate increase in infectious bacteria produced in the ISG15-KO background (2.5 fold for KO-C2 and 1.7 fold for KO-C4) compared to wild-type cells (Fig. 2D). Consistently, we observed a small increase in the bacterial load upon silencing of *ISG15* in HeLa cells (1.3 fold) and in MEFs KO for *ISG15* (1.3 fold Fig. S3, left panels). The ability for the bacteria to attach to (Fig. S3, middle panels) and to penetrate into (Fig. S3, right panels) cells were not dependent on ISG15 expression in the two cellular backgrounds, indicating that the increase in bacterial load is linked to an increase in replication capacity, not in higher susceptibility to bacterial invasion. We concluded from these experiments that ISG15 expression restricted *C. trachomatis* development. The effect was however moderate and did probably not account for the strong increase in inflammatory cytokine production observed in the absence of ISG15 expression, as analysis of cytokine production at the single cell level showed only a weak correlation between bacterial load and cytokine production (Fig. S4).

#### IFN-I signaling does not account for ISG15 upregulation upon Chlamydia infection

We next investigated the signaling pathway(s) leading to the transcriptional up-regulation of *ISG15* expression during *C. trachomatis* infection. We first asked how specific the effect was on *ISG15* transcription as opposed to other ISGs. As expected, transcripts levels of four commonly studied ISGs, i.e. *IFIT3*, *RIG1*, *ISG56* and *MXA*, were upregulated by IFN-I. In the ISG15-KO HeLa cells IFN-I stimulation resulted in a stronger transcriptional up-regulation of these four genes compared to that observed in control cells (Fig. 3A), a phenomenon already documented (4). The transcription of these *ISGs* were also stimulated by *C. trachomatis* infection, but to a lesser extent than that of *ISG15*, and their induction was dependent on *ISG15* being present (Fig. 3B). This data indicated that, among ISGs, *ISG15* was particularly sensitive to the infectious context. Still, activation of several ISGs was consistent with the hypothesis that *ISG15* stimulation involved autocrine stimulation of cells by INF-I. To test this hypothesis we used U5A epithelial cells. These cells do not respond to IFN-I due to a mutation in the IFNα/β receptor (34). The up-regulation in *ISG15* transcription in U5A cells upon *C. trachomatis* infection could not be distinguished from that in the parental 2fTGH cells (Fig. 3C). This result demonstrated that autocrine secretion of IFN-I did not account for the increase in *ISG15* transcription upon *Chlamydia* infection.

**Fig 3.**
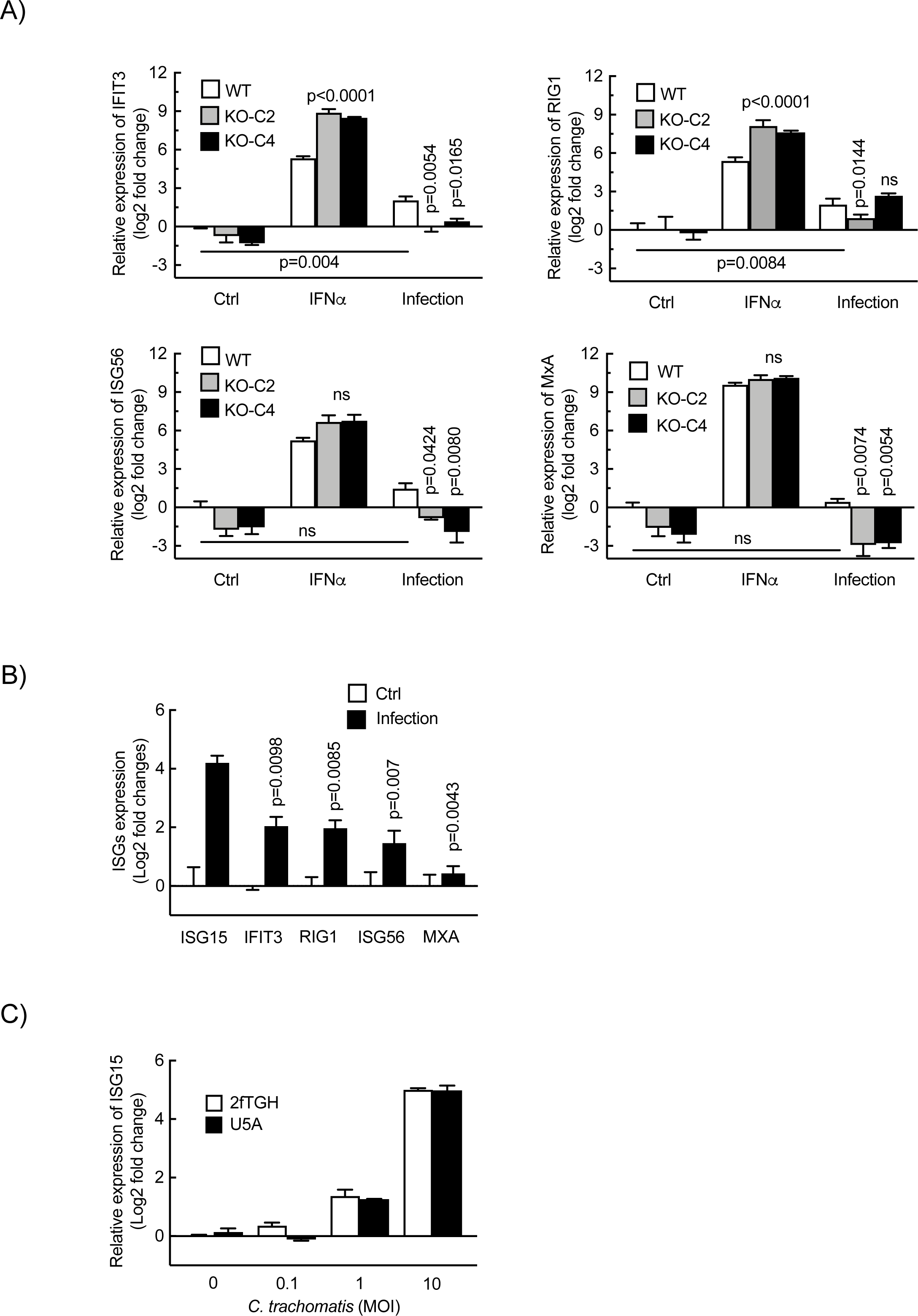
Autocrine IFN signaling is not implicated in *Chlamydia*-induced ISG15 expression by epithelial cells. A) Two clones of ISG15-KO HeLa cells and wild-type cells were infected with *C. trachomatis* (MOI = 1) for 42 h or incubated with IFNα (10 ng/ml) for 24 h. After treatment, the transcript levels of the ISGs including *IFIT3*, *RIG1*, *ISG56* and *MXA* normalized to *actin* were determined and expressed relative to non-infected WT cells. B) HeLa cells were infected with *C. trachomatis* (MOI = 1) for 42 h. The expression of *ISGs* were normalized to actin and are expressed relative to non-infected cells. C) U5A epithelial cells with mutation of IFN-I receptor and the parental 2fTGH cells were infected by *C. trachomatis* at indicated MOI and *ISG15* transcript level normalized to *actin* was determined 42 hpi. The data represent 3 independent experiments. Unpaired t test in A and one-way ANOVA test in B were performed. The p-values of the comparison with WT cells (A), *Chlamydia*-induced ISG15 (B), or as indicated are shown.

#### ISG15 upregulation is not mediated by the PI3K/Akt and NF-κB signaling pathways

Given that *C. trachomatis* infection activates the PI3K/Akt signaling pathway(35), we then tested its implication in the upregulation of ISG15. Surprisingly, only low levels of Akt phosphorylation were detected upon *C. trachomatis* infection (Fig. S5A). Wortmannin, a PI3K inhibitor, had no apparent effect on *Chlamydia*-induced ISG15 expression (Fig. S5B). These data indicated that the PI3K/Akt signaling pathway was not implicated.

To test the contribution of the transcription factor NF-κB we used HeLa cells stably expressing p65, a subunit of the most common p50/p65 heterodimer of NF-κB, as a GFP fusion protein under the control of the EF-1α promoter(36). TNFα stimulation was used as a positive control in these experiments. Treatment with TNFα triggered the translocation of p65-GFP to the nucleus but *C. trachomatis* infection did not (Fig. S5C). Gel shift assays also demonstrated that NF-κB was not activated in HeLa cells upon *Chlamydia* infection (Fig. S5D). These results indicated that the transcription factor NF-κB was not implicated in the upregulation of *ISG15* transcription upon *C. trachomatis* infection.

#### ISG15 expression upon infection is triggered by activation of a cGAS/TBK1/STING/IRF3 pathway

We next screened several transcriptional factors or mediators of various signaling pathways for their contribution to *ISG15* up-regulation, i.e. Myd88, RelA, LRRFIP1, RIG1, IRF3, MDA5, TBK1, STING, STAT1 and IRF7. Two days after silencing, the cells were infected with *C. trachomatis*. Cell lysates were collected 42 h later to measure *ISG15* transcripts and protein levels. *IRF3*, *TBK1* and *STING* were the only three genes whose silencing markedly decreased *Chlamydia*-induced *ISG15* transcription in HeLa cells (Fig. 4A). Consistently, their silencing also prevented the increase in ISG15 levels measured by western blot (Fig. 4B). These three genes participate to the same signaling pathways, as STING (stimulator of interferon genes) and the kinase TBK1 interact with each other, and are essential for the activation of IFN-I genes through subsequent activation of the transcription factor IRF3 (37, 38). To confirm their implication in ISG15 upregulation, we tested the effect of the STING activator, c-di-AMP, on ISG15 synthesis. c-di-AMP (BioLog Life Sciences, #C088-01) had no effect on ISG15 expression by HeLa cells when applied in the culture medium, even at concentration of 10 μM (Fig. 4C). In contrast, introducing c-di-AMP into HeLa cells using transfection reagents stimulated ISG15 expression (Fig. 4D). STING can also be activated by cyclic GMP-AMP, a unique secondary messenger synthesized from GTP and ATP by the cytosolic GMP-AMP synthase (cGAS) (39). To test the role of cGAS in ISG15 induction upon *Chlamydia* infection, we silenced the expression of this enzyme in HeLa cells prior to infection. We observed a significant decrease in the amount of *ISG15* transcripts and ISG15 protein (Fig. 4E), indicating that ISG15 is produced in response to cytosolic DNA detection by the cGAS/STING pathway. We also observed an increase in the transcription of *IL6* and *IL8* genes upon cGAS silencing (Fig. 4F). This result is fully consistent with a need for cGAS to induce ISG15 expression during infection, and thereby dampen the expression of inflammatory genes. Thus, these experiments position *ISG15* transcription downstream of the activation of the c-di-AMP/TBK1/STING/IRF3 and cGAS/TBK1/STING/IRF3 pathways upon *C. trachomatis* infection.

**Fig 4.**
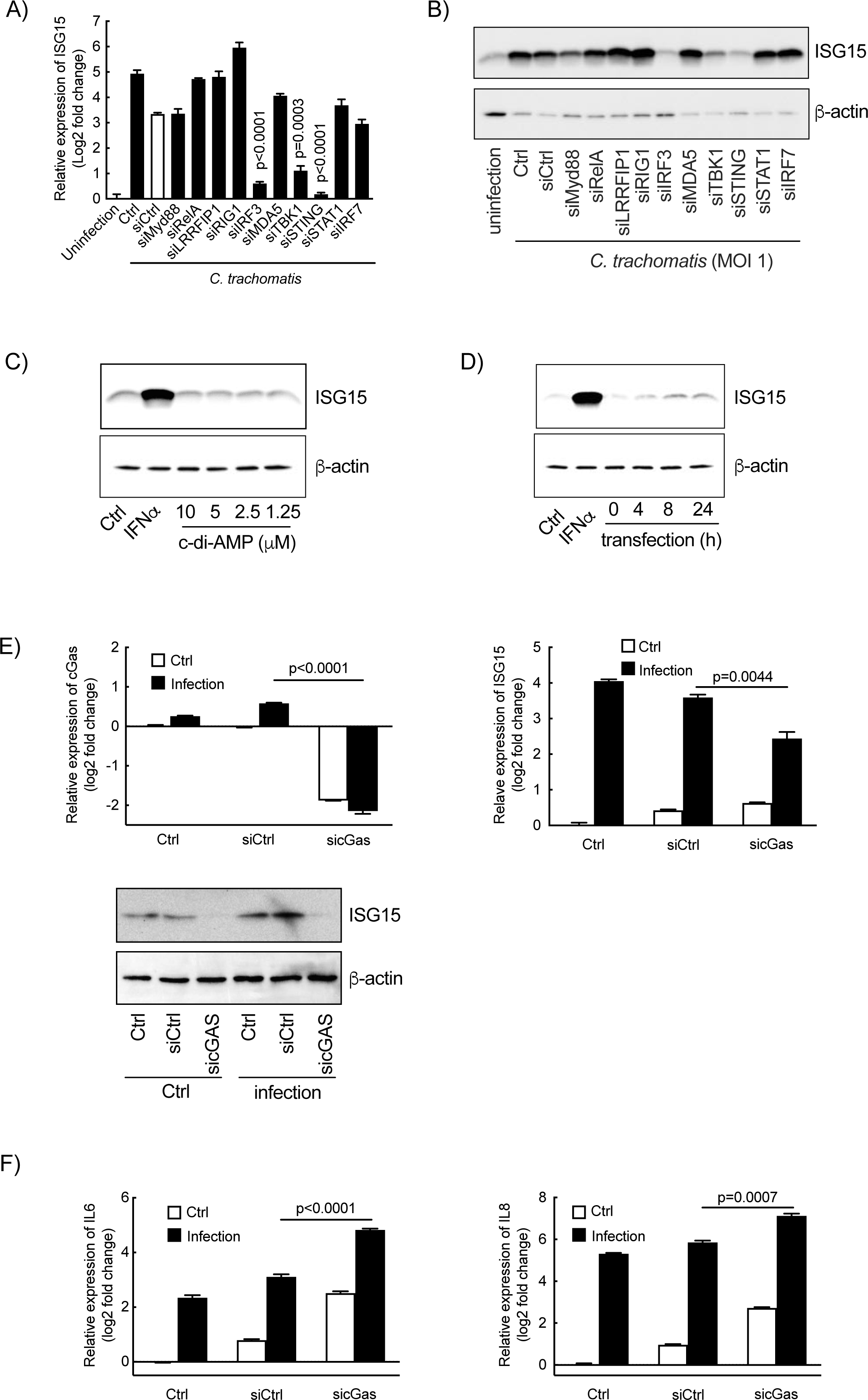
The TBK1/Sting/IRF3 pathway lies upstream of ISG15 synthesis by epithelial cells in response to *Chlamydia* infection. A-B) The indicated genes were silenced by siRNA (30 nM) for 48 h prior to *Chlamydia* infection. Forty-two hours post infection, ISG15 expression was examined either by RT-qPCR (A) or by immunoblot (B). C) HeLa cells were incubated with c-di-AMP at the indicated concentration for 24 h before quantification of the ISG15 levels in whole cell lysates by western blot. D) c-di-AMP (10 μM) was introduced into HeLa cells using transfection reagent at 37 °C for the indicated times. The cells were then washed and were incubated for an additional 24 h before examining ISG15 level in whole cell lysates by western blot. Incubation of HeLa cells with IFNα (10 ng/ml) for 24 h was used as positive control (C & D). E-F) siRNA against *cGAS* or irrelevant oligonucleotides were transfected into HeLa cells for 48 h prior to *Chlamydia* infection. Forty-two hours post infection, the expression of *cGAS* (E, left panel), *ISG15* (E, middle and bottom panels), *IL6* and *IL8* (F) and *actin* were measured. All data represent three independent experiments. The p-values of unpaired t test with the condition “control siRNA” (A) or between the indicated conditions (E, F) are shown.

#### The ISG15 brake on inflammation during Chlamydia infection acts intracellularly independently of ISGylation

ISG15 is a cytoplasmic molecule that is also secreted by multiple cell types, including neutrophils, epithelial cells, monocytes and T cells (6, 40). The extracellular form exerts biological functions, and its receptor has been identified (3). We thus tested the possibility that *Chlamydia*-induced ISG15 was secreted extracellularly and controlled the expression of proinflammatory cytokines by HeLa cells upon bacterial infection from this location. First, the culture medium of infected HeLa cells was collected to probe for the presence of secreted ISG15 by dot blot. Assays using serial quantities of recombinant human ISG15 (rhISG15, R&D systems) determined that the sensitivity of the technique was sufficient to detect up to 25 ng/ml rhISG15 (Fig. 5A). No signal was detected when the membrane was probed with culture medium from infected wells (Fig. 5A), indicating that extracellular ISG15 concentration is lower than this threshold. Furthermore, incubation of ISG15-KO HeLa cells with up to 1 μg/ml rhISG15 had no effect on the basal, nor on the infection-induced, levels of *IL6* and *IL8* transcripts (Fig. 5B). These results demonstrate that extracellular ISG15 does not affect *Chlamydia*-induced inflammation, indicating that it exerts this activity inside the cell.

**Fig 5.**
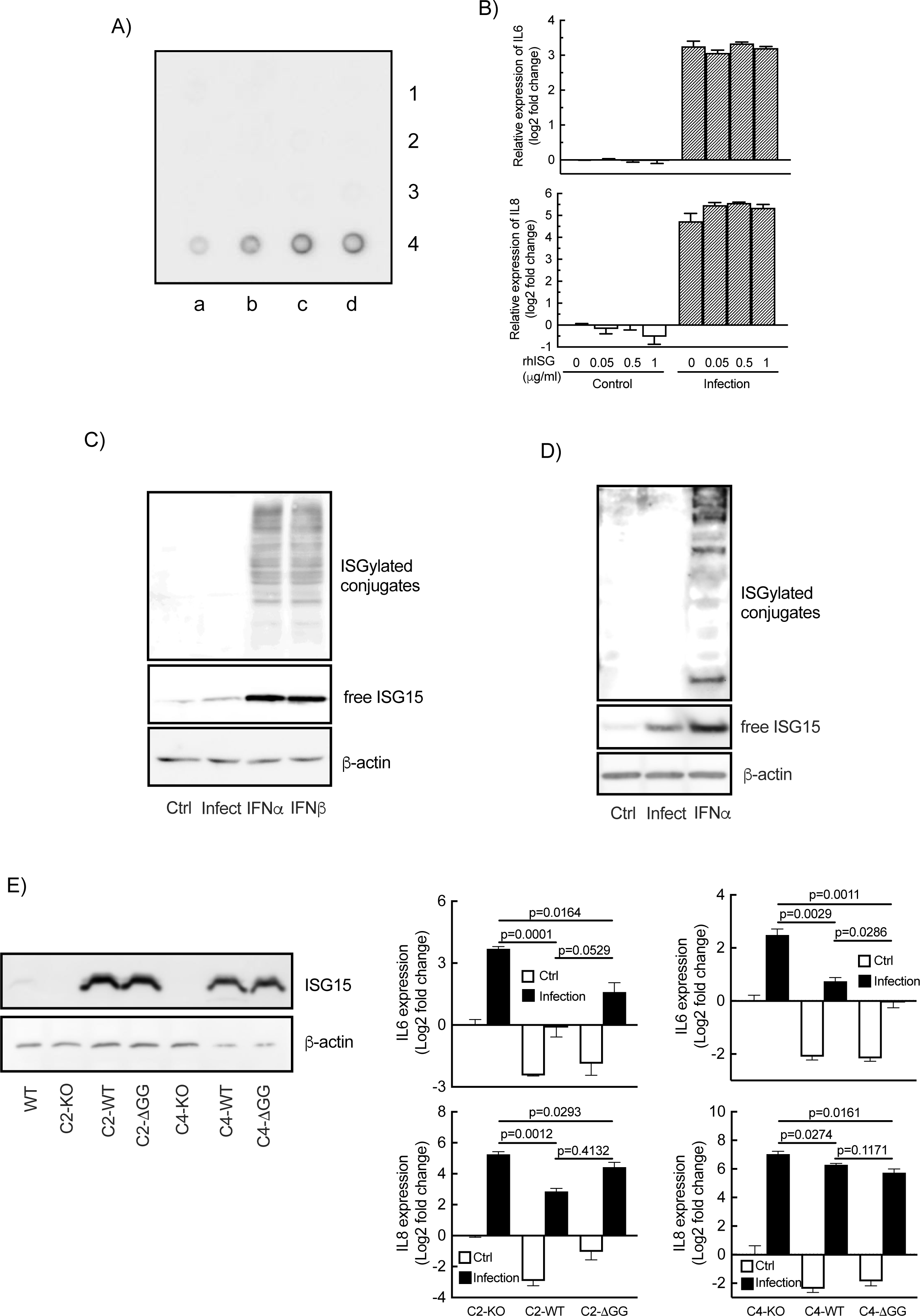
ISG15 regulates inflammation from an intracellular location and independently of ISGylation. A) Culture medium was collected at different times post infection for ISG15 detection by dot-blot. First row: 0, 3, 6, 9 hpi; 2^nd^ row: 12, 18, 24 and 30 hpi; 3^rd^ row: 36, 42 and 48 hpi; 4^th^ row: 25, 50, 100 and 200 ng/ml of rhISG15. B) ISG15-KO HeLa cells were pre-treated with the indicated concentration of rhISG15 for 30 min prior to *Chlamydia* infection (MOI=1). Cells were further incubated for 42 h in the presence of rhISG15 before RNA extraction and quantification of *IL6* and *IL8* expression by RT-qPCR. C) HeLa cells were incubated with *C. trachomatis* (MOI=1) for 42 h or with IFN-I (10 ng/ml) for 24 h, lysed, and the amount of ISG15 (free form & conjugates) was analyzed by immunoblot on whole cell lysates. D) Primary cervical epithelial cells were treated with IFNα (10 ng/ml) for 24 h or with *C. trachomatis* (MOI=10) for 42 h followed by ISG15 detection. Top panels of C and D show overexposed images of the upper part of the membranes to visualize ISGylated proteins. E) ISG15-KO cells (C2 & C4) or complemented cells (with ISG15 WT or 1′GG) were infected or not with *C. trachomatis* (MOI=1) for 42 h before measuring the transcription of *IL6* and *IL8*. The panel on the left shows ISG15 expression in the complemented cells. The images displayed are representative of 2 independent experiments in A, and of 3 independent experiments in D using primary cells from 3 individuals. All other data correspond to 3 independent experiments. The p-values of unpaired t test between the indicated groups are shown.

Intracellular ISG15 displays biological activities either as a free molecule or through conjugation to other proteins, a process called ISGylation (40, 41). We tested whether *Chlamydia* infection was accompanied with global changes in the level of ISGylation. IFNα and β were used as positive controls. As expected, HeLa cell exposure to these cytokines elicited a strong increase in ISG15 expression and protein ISGylation (Fig. 5C). Infection induced ISG15 expression, but to a lower extent than IFN-I did, and protein ISGylation was below detection limit (Fig. 5C). Similar results were obtained in *Chlamydia*-infected primary cervical epithelial cells of patients (Fig. 5D). To determine whether ISGylation was implicated in the ISG15-mediated restriction on inflammation we complemented two ISG15-KO clones with truncated ISG15-ΔGG, that is not competent for ISGylation (42). We observed that expression of ISG15-ΔGG, attenuated *Chlamydia*-induced expression of IL6 and IL8 to the same extent as ISG15-WT (Fig. 5E). These experiments demonstrate that the ability for ISG15 to dampen inflammation does not involve ISGylation.

#### Lack of ISG15 results in exacerbated tissue damage in a mouse model of infection

We finally tested the consequence of ISG15 deficiency using a trans-cervical mouse infection model (29). In this model, INFγ^-/-^ mice showed decreased *C. trachomatis* clearance on day 6 of infection, indicating that IFNγ was important to restrict bacterial growth. Given the need of ISG15 for a strong IFNγ response in the case of *M. tuberculosis* infection (6), we expected to also observe delayed *C. trachomatis* clearance in the ISG15-KO animals. Surprisingly, bacterial loads measured on day 3 and 6 were not different between the two groups of mice (Fig. 6A). However, 4 weeks post infection, we observed higher bacterial load (Fig. 6A) and higher levels for several proinflammatory cytokines in the genital tract of ISG15-KO animals, i.e. MIP1α, MIP1β, Rantes, IL13 and Eotaxin (Fig. 6B & S6). ISG15-KO animals displayed more severe signs of hydrosalpinx compared to wild type mice (Fig. 6C). Thus, ISG15 deficiency weakened the natural ability of mice to clear *C. trachomatis* infection, and exacerbated inflammation and subsequent tissue damage.

**Fig 6.**
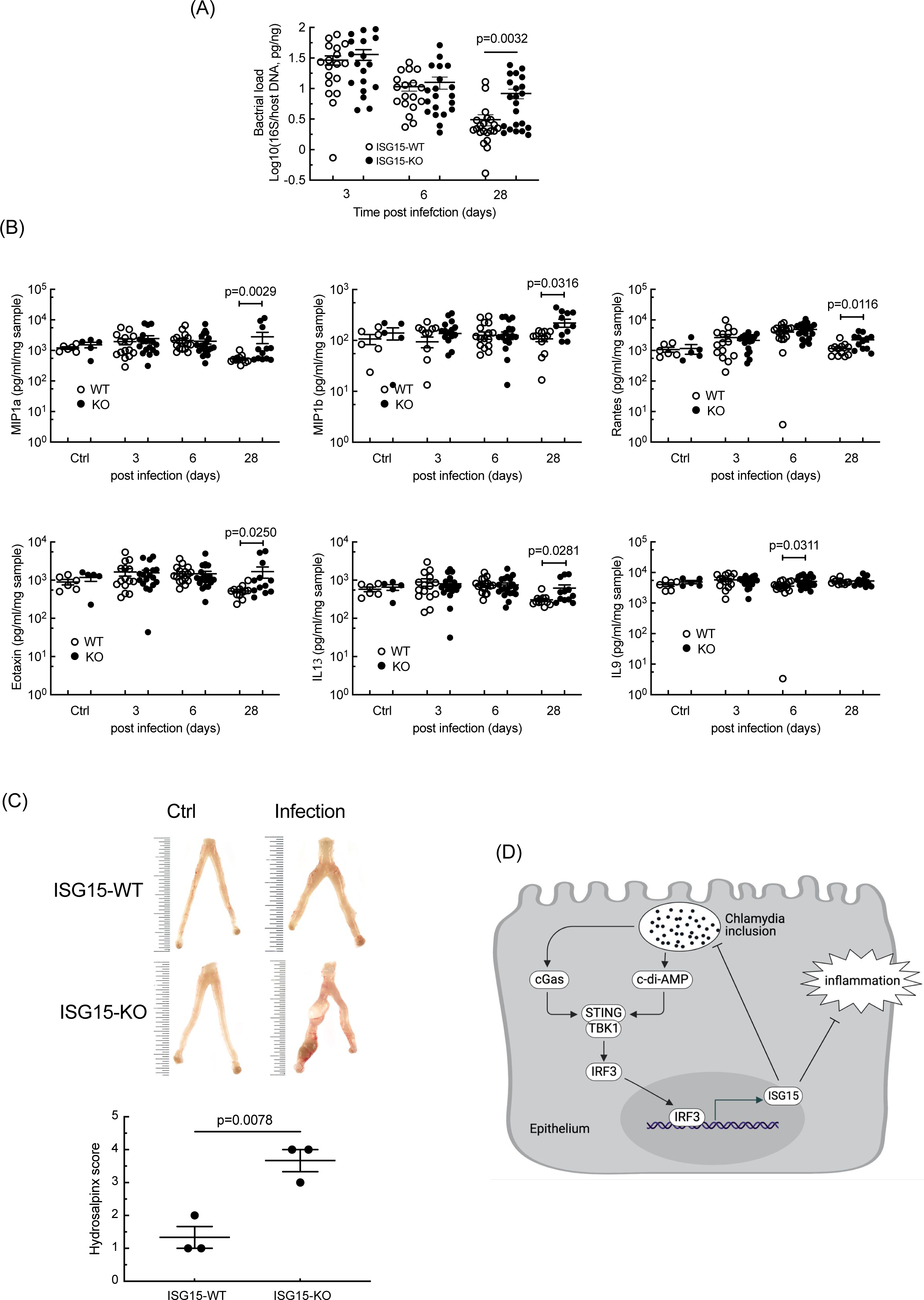
ISG15 limits the inflammatory response in the upper FGT of mice infected with *C. trachomatis*. ISG15-KO and ISG15-WT mice were infected by introducing *C. trachomatis* into the uterine horn trans-cervically. The upper FGTs were collected at the indicated times post infection. Bacterial loads (A) and inflammatory cytokines (B) in FGT homogenates were determined using quantitative PCR and multi-plex ELISA assays, respectively. Each dot represents one animal. C) Representative FGTs (top) and hydrosalpinx index for six infected animals (bottom). The smallest interval of the scale in this image represents 0.05 cm. The p-value of unpaired t test between indicated groups are shown. D) Graphical summary. *Chlamydia* infection of epithelial cells is sensed by cGas, which activates the STING/TBK1/IRF3 signaling cascade. Direct release of c-di-AMP by the bacteria may also be involved. IRF3 activates transcription of *ISG15*. Increase in intracellular free ISG15 restricts bacterial proliferation and the transcription of the host inflammatory cytokines IL6 and IL8, independently of ISGylation.

## Discussion

The lower FGT of women is exposed to the vaginal microbiota, and occasionally to a variety of sexually transmitted pathogens including HPV, HIV and *C. trachomatis*. The latter induce an inflammatory response aiming at the elimination of the bacteria, but which is also at the root of the pathological consequences of *Chlamydia* infection. In this report, we show that infection of primary cervical epithelial cells, as well as of the HeLa cell line, by *C. trachomatis* elicits a strong induction of the expression of the ubiquitin-like molecule ISG15, and that this molecule acts as a brake on the immune response to infection. Reducing or knocking-out the expression of *ISG15*, in different cellular backgrounds, resulted in an increase in the transcription and secretion of two inflammatory cytokines, IL6 and IL8, which was rescued by ISG15 complementation into ISG15-KO cells. Quantification of cytokines and chemokines by ELISA confirmed that ISG15 broadly repressed the immune response to infection. As discussed below, we describe the signaling pathway that triggers ISG15 expression. Knocking-out ISG15 expression in mice delayed *C. trachomatis* clearance and resulted in increased tissue damage. Overall, the induction of ISG15 in epithelial cells upon *C. trachomatis* infection benefits the host by limiting bacterial replication and circumventing the inflammatory response (Fig. 6D). Induction of ISG15 expression upon *C. trachomatis* infection was observed in HeLa cells, in primary cells, in MEFs (this report) and in a Fallopian-tube derived organoid model (43). We showed that silencing the TBK1/STING/IRF3 signaling cascade was sufficient to prevent *ISG15* induction upon *C. trachomatis* infection. During viral infection, activation of the TBK1/STING/IRF3 signaling cascade leads to the expression of IFN-I (37, 38). IFN-I binding to its receptor IFNAR triggers the canonical STAT signaling cascade, resulting in the transcription of many IFN-responsive genes, including *ISG15*. Interestingly, in the infection situation studied here, *ISG15* transcription was induced even in the absence of IFNAR signaling (Fig. 3C). The increase in ISG15 level upon another bacterial infection, *Listeria monocytogenes*, was also independent of IFN-I (7). It is important to note that the increase in ISG15 upon *C. trachomatis* infection was lower than upon IFN-I induction (Fig. 1D-E, 5C-D). Therefore, the amplification of *ISG15* transcription upon IFN-I signaling may mask the intrinsic capacity of the TBK1/STING/IRF3 signaling cascade to directly upregulate *ISG15,* and account for the apparent difference between viral and bacterial infection regarding the regulation of *ISG15* transcription. In other words, the low level of IFN-I induction in epithelial cells infected by *Chlamydia* enables to detect the direct regulation of *ISG15* expression by the TBK1/STING/IRF3 signaling cascade.

Silencing cGAS resulted in a strong decrease in the *Chlamydia*-induced upregulation of ISG15, indicating that detection of nucleic acids was involved, like in viral infection. This is consistent with the observation that c-di-AMP is a prominent ligand for STING-mediated activation of IFN responses during *C. trachomatis* infection(44). Interestingly, *C. trachomatis* possesses an enzyme that synthesizes c-di-AMP (44). This might contribute to direct STING activation, in addition to cGAS generated cyclic GMP-AMP. In contrast, *Chlamydia*-induced IL1β in macrophages was dependent of STING but independent of cGAS (45).

Silencing, or knocking-out, the expression of ISG15 enhanced the expression of the inflammatory cytokines IL6 and IL8, which was rescued by complementation with *ISG15* under a constitutive promoter (Fig. 2C & 5E), indicating that ISG15 acts as a negative regulator of inflammation. Consistent with this hypothesis, the transcriptional induction of *IL6* and *IL8* starts late in HeLa cells, when each vacuole contains already several hundreds of bacteria (46, 47). In contrast, *ISG15* transcription is induced within 6 h of infection. These kinetics are consistent with a scenario in which ISG15 refrains the induction of the pro-inflammatory signals. This phenomenon might be favorable to the bacteria, by delaying the immune alert until close to the completion of the infectious cycle. However, our *in vivo* data support overall a beneficial role for ISG15 in limiting *C. trachomatis* infection and the associated inflammation, since ISG15 deficient mice showed a delayed bacterial clearance and more tissue damage than wild type mice. In contrast, overexpression of human ISG15 enhanced IL6 and IL8 production induced by *L. monocytogenes* infection in HeLa cells (7).

Mechanistically we uncovered another important difference when comparing *Chlamydia* to other bacterial infections. Both *Listeria* and *Mycobaterium* have been shown to elicit ISGylation *in vitro* and *in vivo*, respectively (7, 48). In contrast, in spite of the increase in free ISG15, ISGylated proteins remained below detection threshold by western blot in *C. trachomatis* infected cells (Fig. 5C&D). This was not due to an increase in USP18, a protease reversing ISGylation (49), since the protease level was stable over the course of infection (data not shown). *C. trachomatis* secrete two proteins, one with deubiquitinase and acyltransferase activity, the second with deubiquitinase activity (50, 51). These enzymes did not utilize ISG15 to form conjugates *in vitro*, indicating that they are not responsible for the low level of ISGylation in *C. trachomatis* infected cells (50). Whether another secreted bacterial factor limits ISGylation remains to be explored. In any case, we showed here that ISGylation was dispensable for the anti-inflammatory activity of ISG15 in epithelial cells, and that this activity was exerted intracellularly. The mechanism(s) by which free ISG15 level modulate the transcription of *IL6* and *IL8* require further investigation.

We were surprised to observe no difference in bacterial load at 3 and 6 days of infection between wild-type and ISG15 KO mice (Fig. 6A). Based on results obtained in *Mycobacterium*-infected patients and mice with ISG15 deficiency (6), we expected the ISG15 KO mice to show an impaired IFN-γ response. However, similar levels of IFN-γ were detected in ISG15-WT and ISG15-KO mice (Fig. S6). These data suggest that in the context of *C. trachomatis* infection ISG15 only plays a marginal role, if any, on regulating IFN-γ levels.

Finally, this work showed that ISG15 had a beneficial role in a mouse model of *C. trachomatis* infection, as its absence resulted in delayed bacterial clearance and increased tissue damage. We showed that its activity on limiting the immune response of epithelial cells was exerted as a free intracellular molecule. However, it is possible that other modes of actions contribute to harnessing the immune response in the animal, with different modes of action depending on the cell types.

## Acknowledgements

ISG15 antibody, smartpools of siRNA and GFP-p65 cells were kindly provided by Zhi Li and Lilliana Radoshevich and Chak Hon Luk, respectively. We thank the Service de Chirurgie gynécologie (Institut Mutualiste Montsouris) and the Centre de Recherche Translationnelle (Institut Pasteur) for their contributions. This work was supported by an ERC Grant (NUChLEAR N°282046), the Agence Nationale de la Recherche [TheraEpi - ANR-20-PAMR-0011], the Institut Pasteur and the Centre National de la Recherche Scientifique (CNRS).

---

Fig. S1 Characterization of ISG15-KO cells A) ISG15-KO clones and ISG15-WT cells were incubated with IFN-I (IFNα 10 ng/ml) for 24 h or infected with *C. trachomatis* for 42 h before measuring ISG15 levels on whole cell lysates by western blot. B) ISG15-KO and ISG15-WT cells were infected with *C. trachomatis* for 42 h. After infection, the transcripts of pro-inflammatory cytokines *IL6* and *IL8* were examined. Unpair t test was performed to compare the indicated clones with non-infected and infected WT cells, respectively, and the p-values are shown. C) ISG15-KO C2 and C4 cells were complemented for *ISG15* expression. After puromycin selection, cellular pools were imagined in the green channel, as GFP is co-expressed with ISG15. The results are representative of 2 independent experiments.

Fig. S2 ISG15 dampens the host immune response to *C. trachomatis* infection. *ISG15*-KO Hela cells (clone C2) complemented or not with ISG15 was infected with or not with *C. trachomatis* (MOI=1). Classical actors of the immune response were quantified by ELISA in the culture supernatant 42 hpi. Panel (A) displays the extracellular levels of the proteins induced by infection in an ISG15-sensitive manner, panel (B) displays the extracellular levels of the proteins tested that did not follow this pattern. One-way ANOVA was performed for statistical analysis.

Fig. S3 ISG15 depletion favors bacterial growth. A) HeLa cells transfected with siISG15 or irrelevant oligonucleotides were infected with *C. trachomatis* for 42 h followed by the quantification of bacteria loads using qPCR (left panel). For bacterial binding study, pre-chilled siISG15-treated or non-treated HeLa cells were incubated with *Chlamydia* LGV-L2^IncD^-GFP (MOI=30) at 4 °C for 4 h. After washing, cells were fixed, stained nuclear DNA and attached GFP-bacteria were quantified under Deltavision™ microscope (middle panel). For bacterial entry study, siRNA-treated cells seeded on coverslip were infected for 45 min at 4°C. After washes, the cells were incubated at 37 °C for the indicated times before staining extracellular bacteria and bacterial entry was analyzed as described in the methodology section (right panel). B) The bacterial binding and entry studies were performed as that in (A) in ISG15-deficient and WT MEFs. Three independent experiments for the studies of bacterial load were performed. The binding and entry studies are representative of two independent experiments. Each dot in the right panels represents quantification of one field. Unpaired t-test was conducted to compare bacterial burdens.

Fig. S4 Single cell analysis of bacterial load and cytokine production. HeLa cells were infected or not with LGV-L2^IncD^GFP bacteria (MOI=1) for 30 h (in the presence of brefeldin A for the last 6 h) before fixation and staining with PC7-conjugated anti-IL6 (upper panels) or PE-conjugated anti-IL8 (lower panels) antibodies. The LGV^+^/IL6^+^ or LGV^+^/IL8^+^ population (red rectangles) was further analyzed to compare bacterial load and cytokine level. Pearson’s correlation coefficient (r) and P-value were calculated with Prism 9 with a two-tailed P-value.

Fig. S5 PI3K/Akt and NF-κB signaling pathways are not involved in *Chlamydia*-induced ISG15 synthesis by epithelial cells. A) HeLa cells were incubated with *C. trachomatis* (MOI = 1) and Akt phosphorylation was examined by western blot at the indicated times post infection. B) HeLa cells were pre-treated with wortmannin for 30 min prior to *Chlamydia* infection. ISG15 expression was determined by western blot 40 hpi. C) HeLa cells constitutively expressing p65-GFP fusion protein under the control of the EF-1α promoter were incubated with TNFα (10 ng/ml) for 15 min or with *Chlamydia* (MOI=1) for the indicated time, prior to fixation and immunofluorescence analysis to monitor the nuclear translocation of p65. D) HeLa cells were infected with *C. trachomatis* (MOI = 1). Twenty-four hours (upper panel) or at the indicated times (lower panel) post infection, the cells were collected for gel shift assay to measure NF-κB activation. TNFα (10 ng/ml) was used as a positive control. The data represent two independent experiments.

Fig. S6 Absence of ISG15 delays *C. trachomatis* clearance and exacerbates tissue damage. ISG15-KO and wild-type mice were infected with *C. trachomatis* by introducing bacteria into the uterine horn. At the indicated times post infection, animals were sacrificed to harvest the upper FGT. The pro-inflammatory cytokines/chemokines in the upper FGT were determined by multi-plex ELISA. Each dot represents one animal. Unpaired t-tests between animal groups at each time point were performed.

**Table S1.**
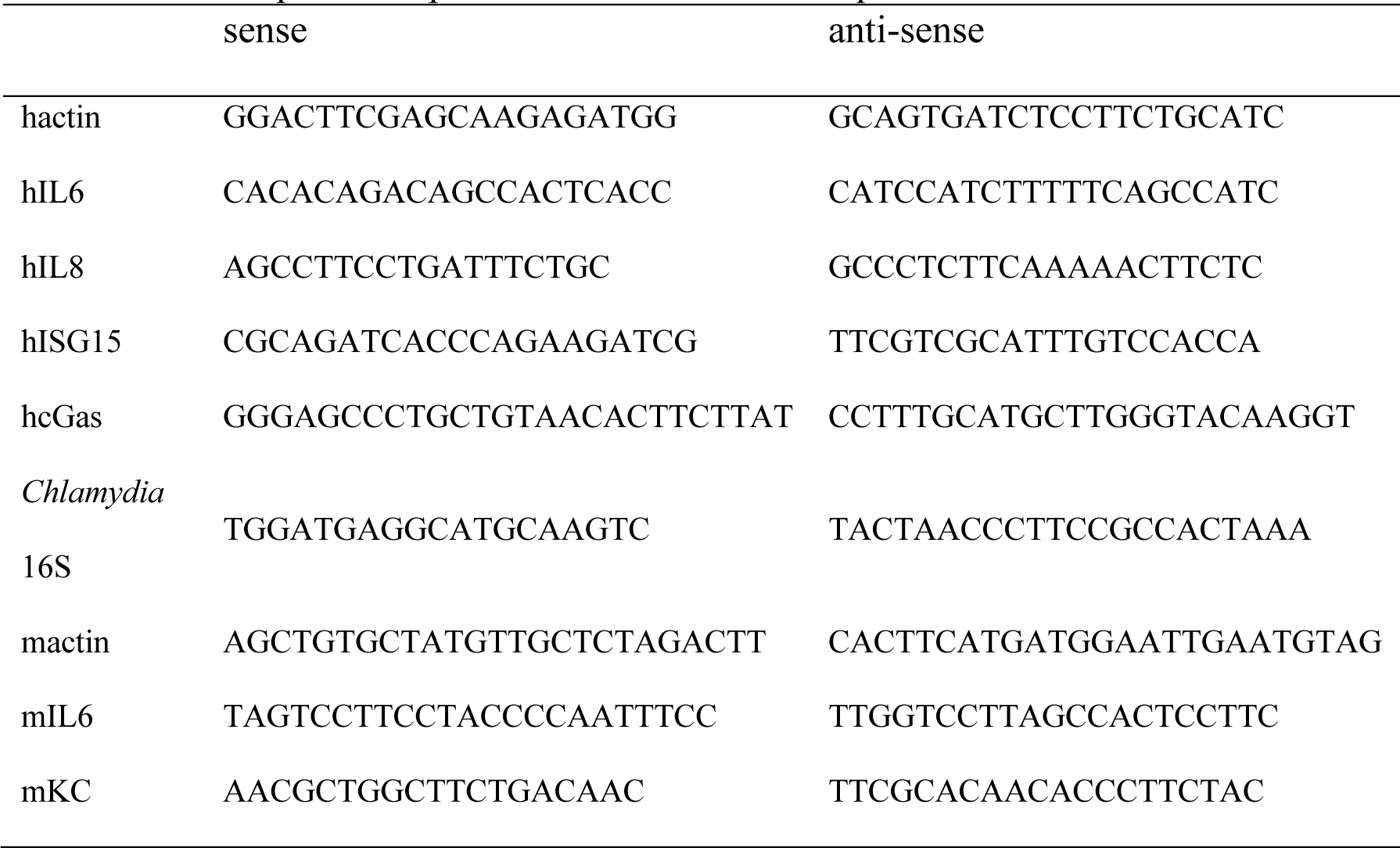
The sequence of primers used for real-time quantitative PCR sense anti-sense.

