## Supplementary figures and images for "*Chlamydia*-driven ISG15 expression dampens the immune response of epithelial cells independently of ISGylation"

### supplementary data

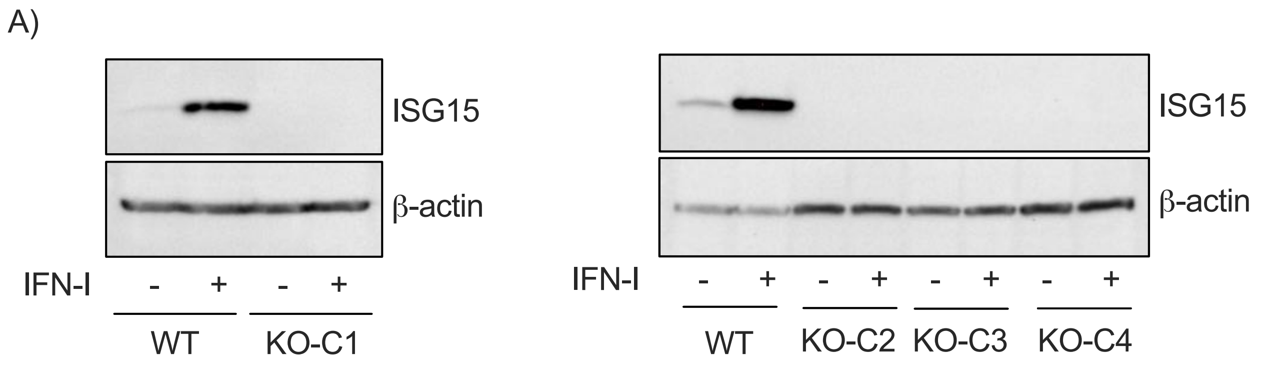

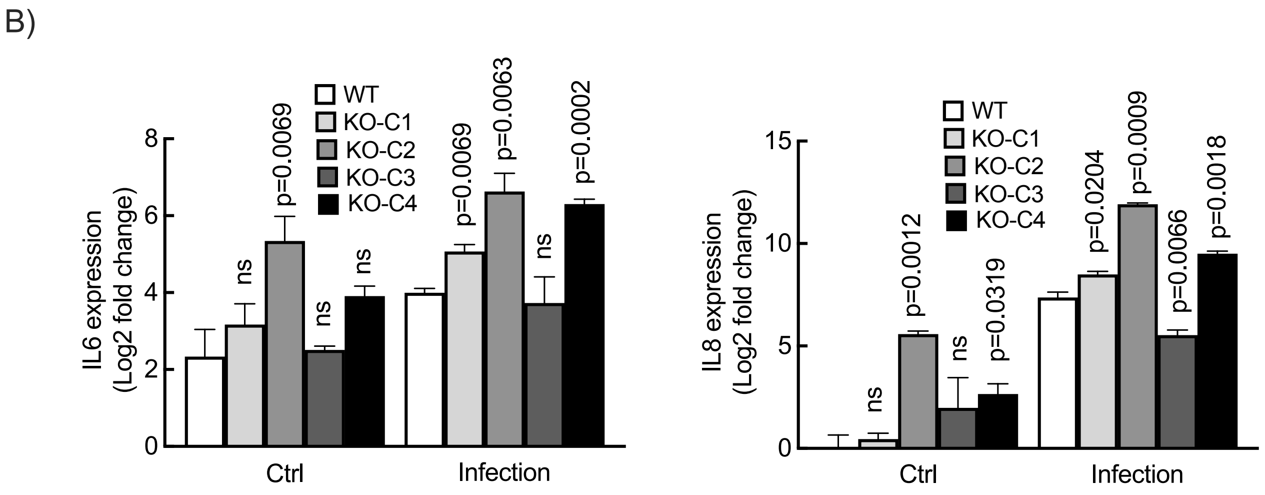

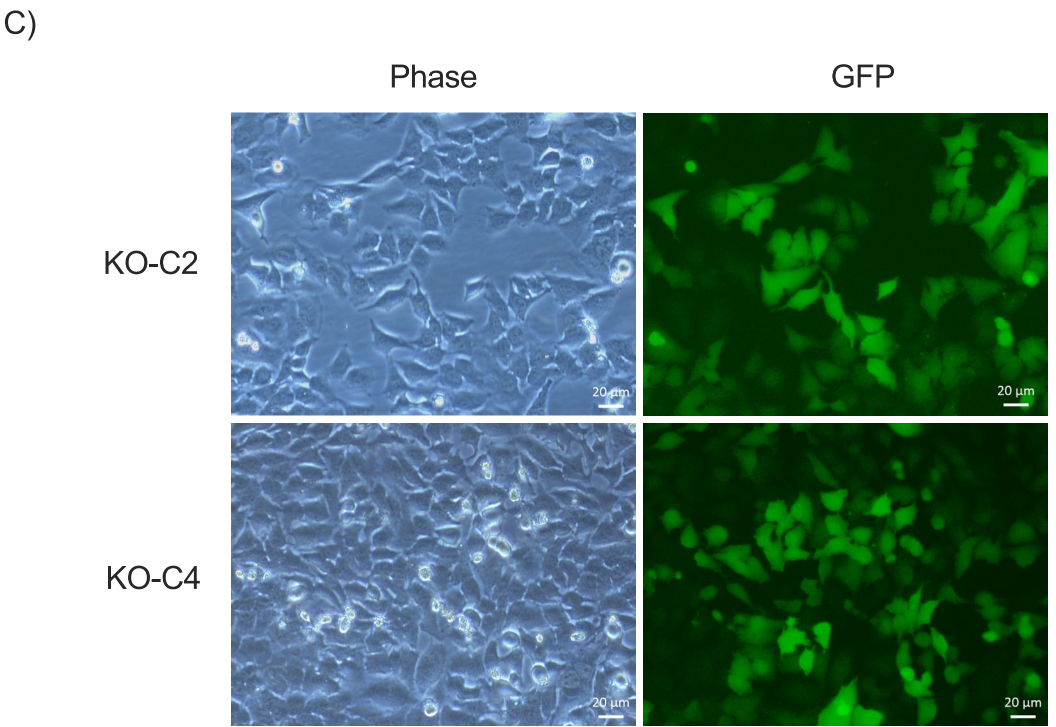

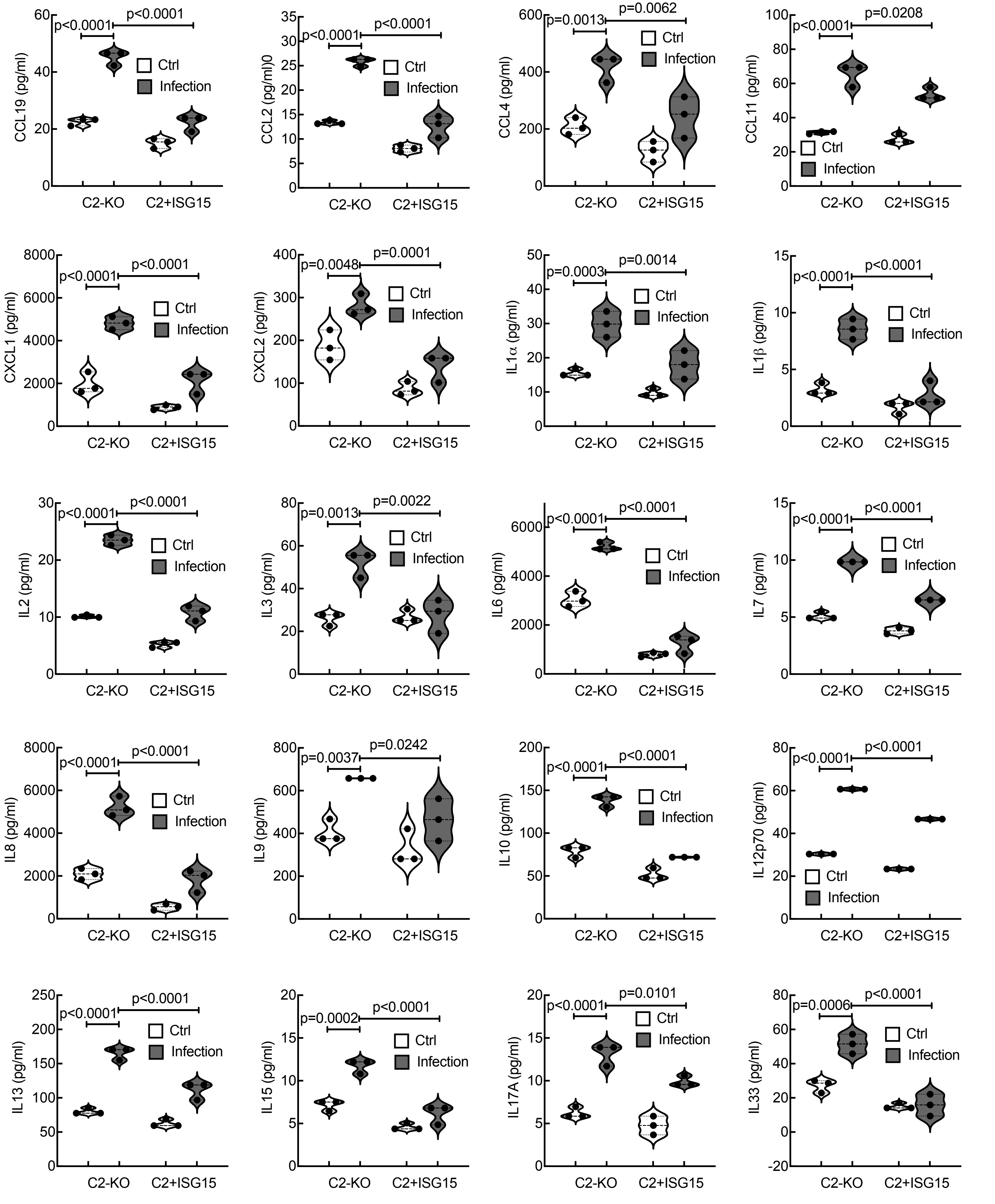

**Fig. S2. (A) Continue**

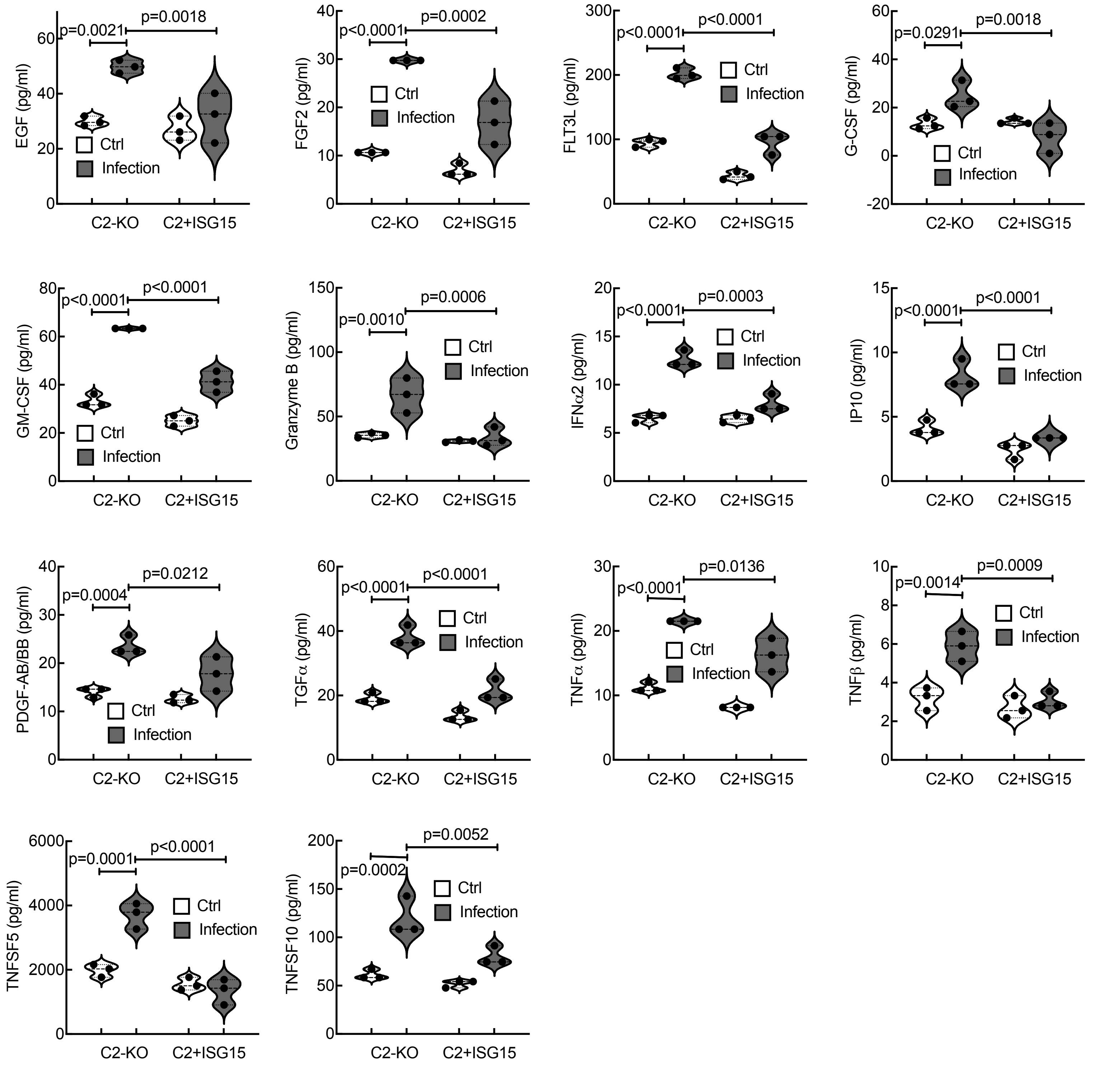

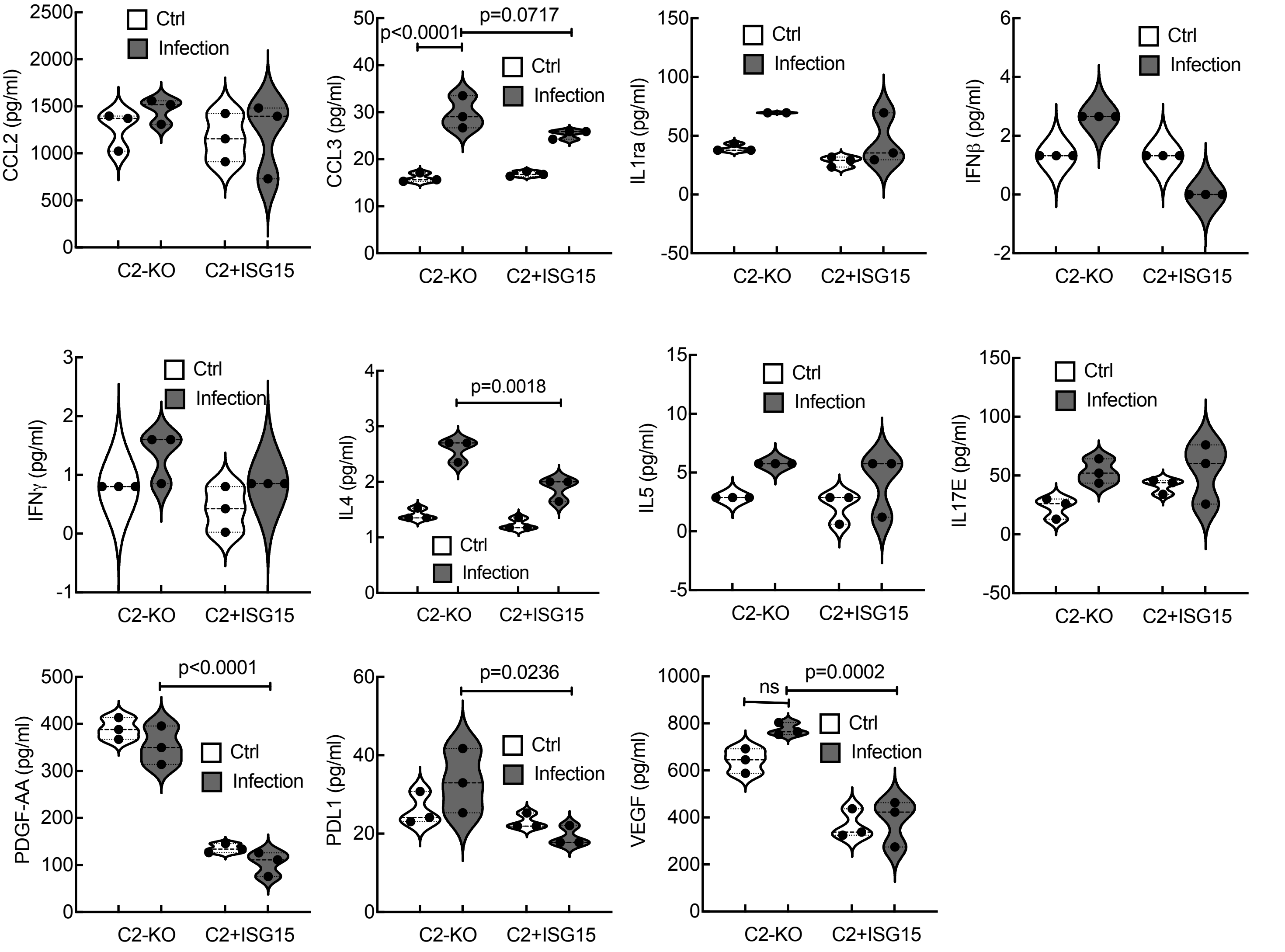

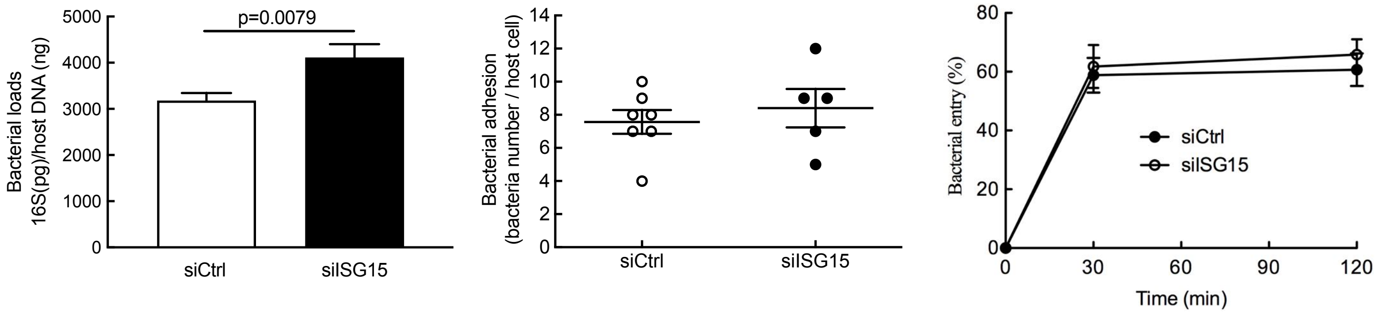

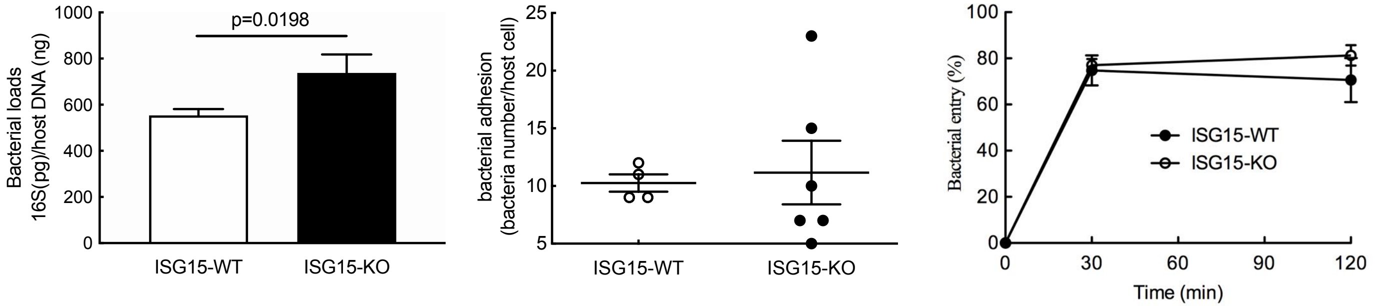

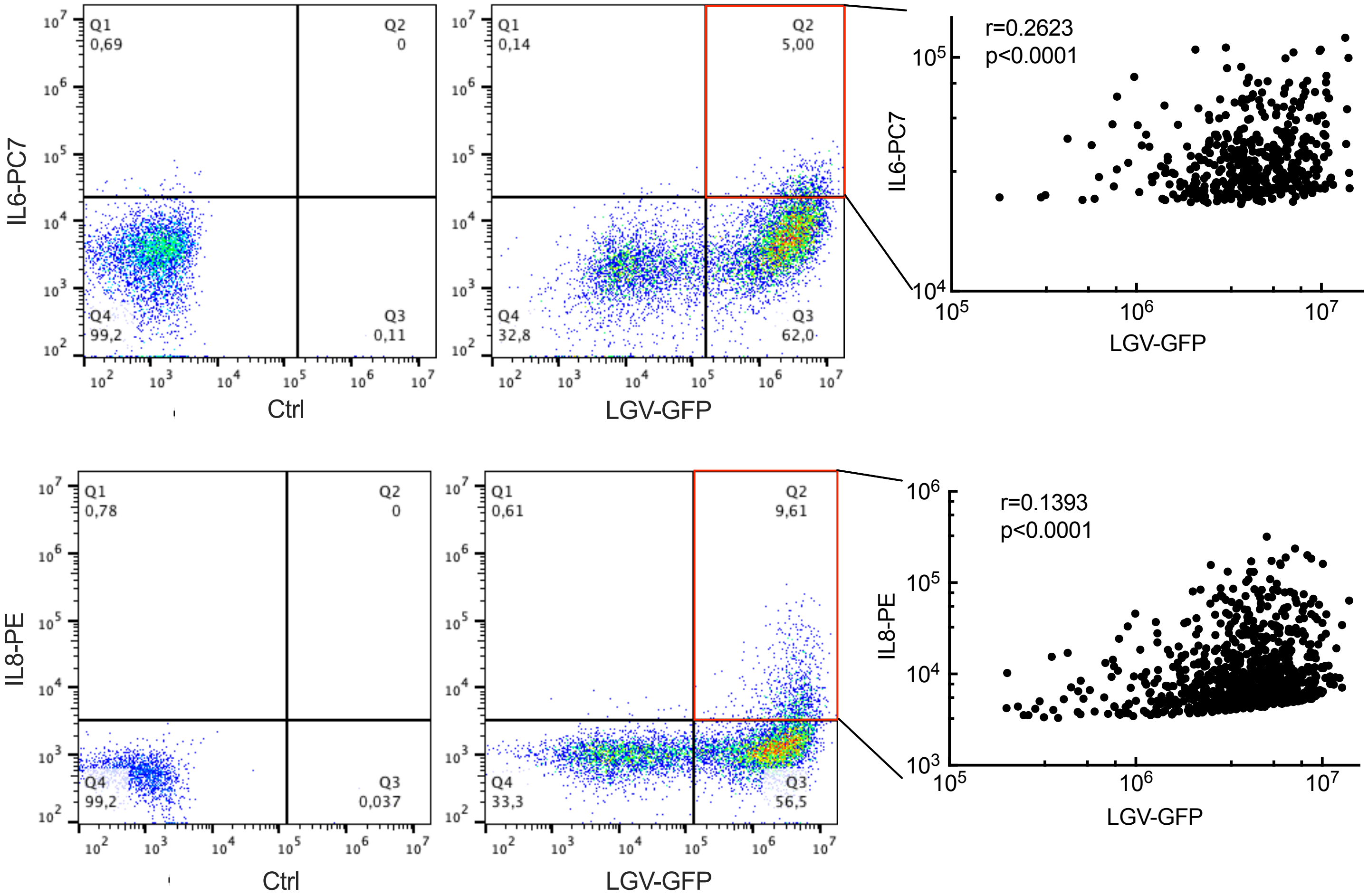

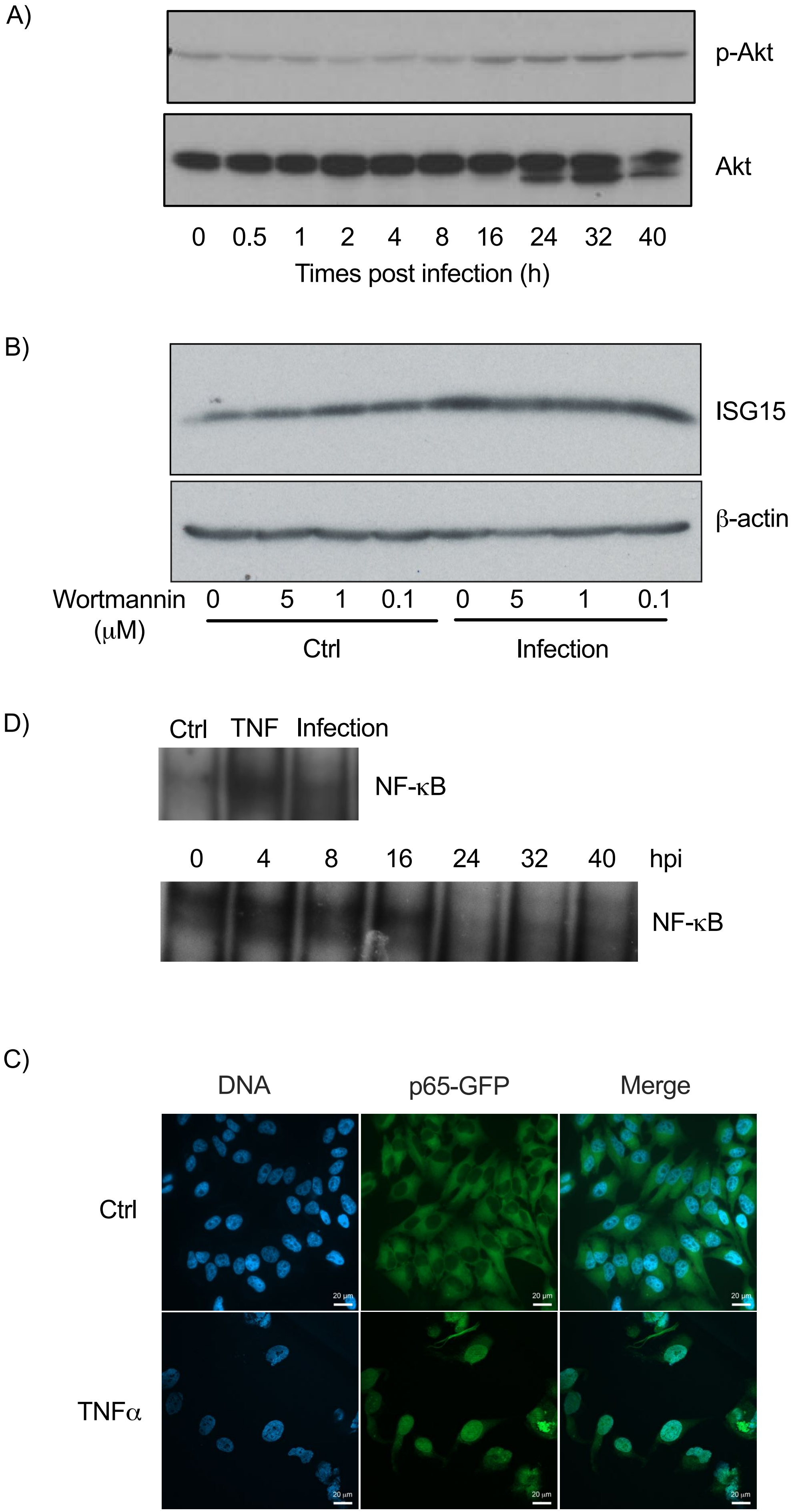

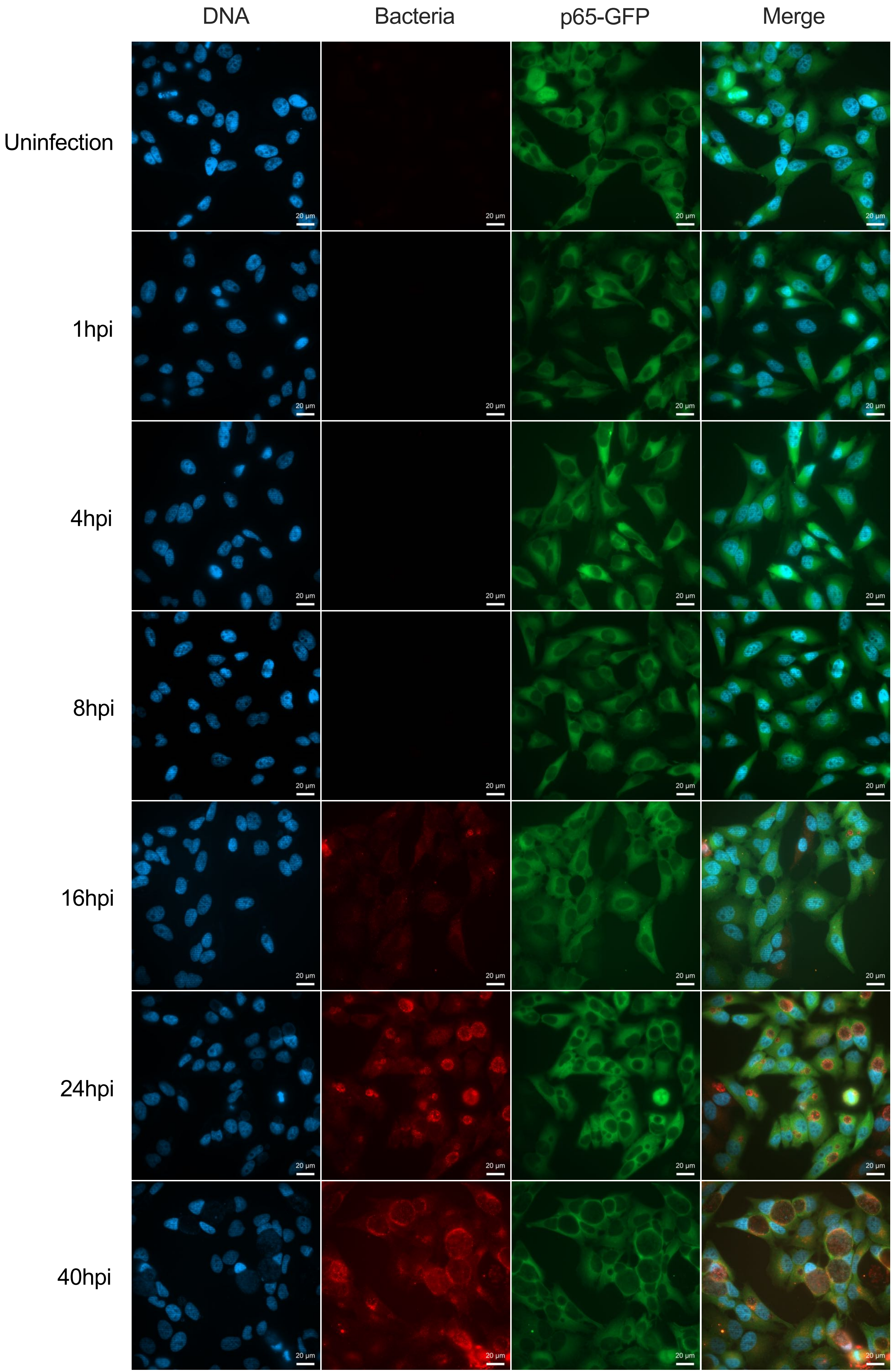

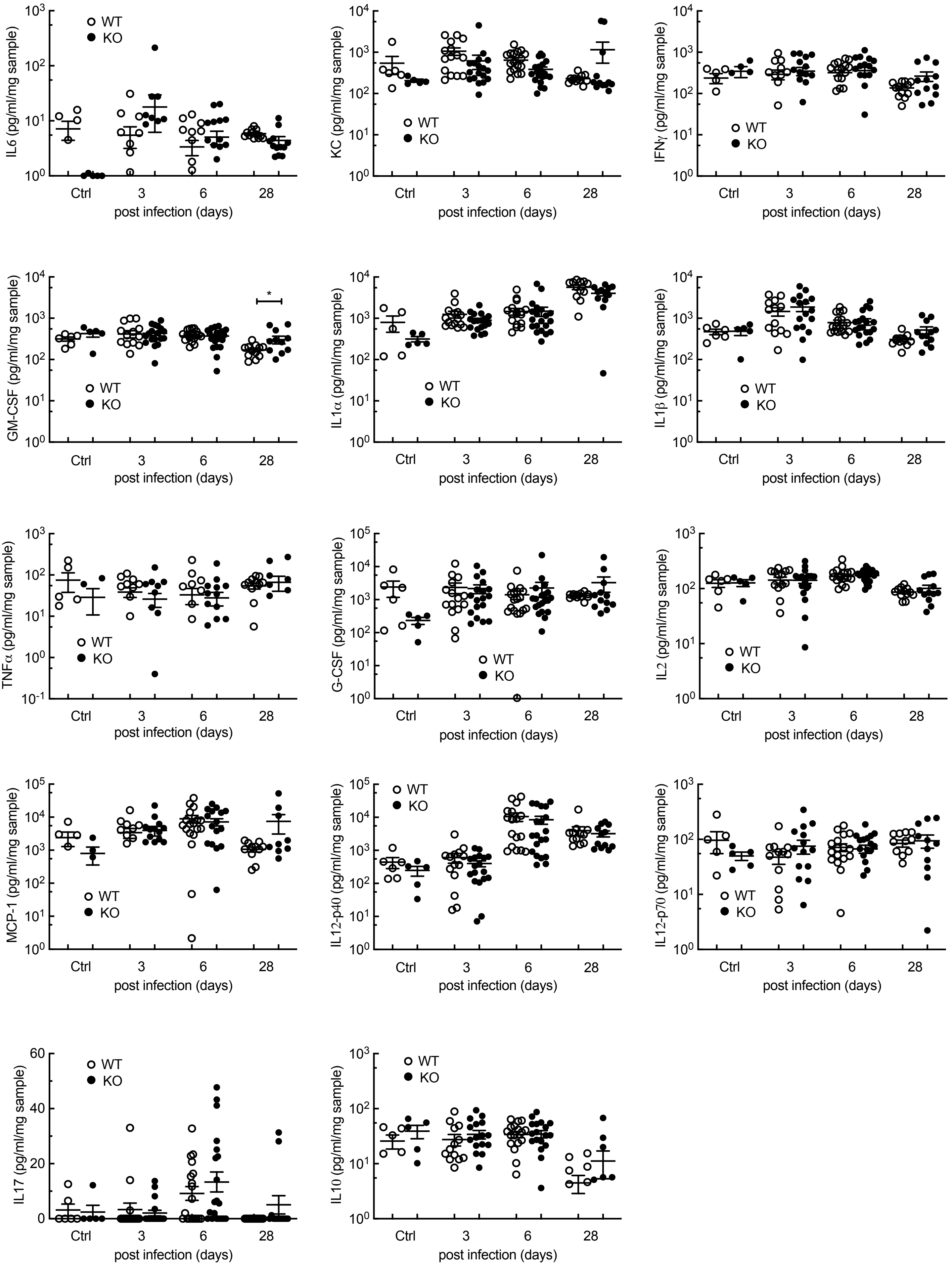
